# Glucose Metabolism Mediates Feedback Control of Innate Immune Signaling in Human Macrophages

**DOI:** 10.64898/2026.08.12.744460

**Authors:** Adebola A. Owolabi, Yetunde I. Kayode, Deanna C. Clemmer, Glenn E. Simmons, Harry E. Taylor

**Affiliations:** Department of Microbiology & Immunology, SUNY Upstate Medical University, Syracuse, NY 13210, USA; Department of Biomedical Sciences, Cornell University College of Veterinary Medicine, Ithaca, NY 14853-64014, USA

## Abstract

Glucose metabolism is pivotal in regulating innate immune responses in primary human monocyte-derived macrophages (MDMs). Lipopolysaccharide (LPS) stimulation induces both inflammatory and antiviral programs; however, despite the established importance of glucose metabolism in these responses, its precise role in coordinating them remains poorly defined. Here, we identify the STAT1/NF-κB/IRF5 signaling axis as a key mediator linking glucose metabolism to inflammatory responses through the upregulation of the rate-limiting glycolytic enzyme PFKFB3. We found that LPS triggered delayed expression and activation of NF-κB p65, accompanied by increased expression of inflammatory target genes, including CD38 and CD40. Using complementary pharmacological and genetic approaches, we demonstrate that glycolysis and PFKFB3 activity are required for NF-κB p65 expression and activation. Strikingly, inhibition of PFKFB3 also suppressed LPS-induced STAT1 activation and nuclear translocation, revealing a glucose-dependent amplification loop that potentiates STAT1-mediated antiviral and NF-κB p65-mediated inflammatory responses. Collectively, these findings establish a mechanistic link between glycolytic metabolism and STAT1/IRF5- and NF-κB-dependent transcriptional programs in human MDMs responding to LPS, highlighting potential therapeutic targets for modulating innate immune responses in inflammatory disease.

## INTRODUCTION

Innate immunity represents the body’s first line of defense against pathogens.^1–3^ It is coordinated by cells of common myeloid origin, including monocytes, macrophages, dendritic cells, and neutrophils,^4, 5^ through the activation of downstream transcription factors such as Nuclear Factor kappa B (NF-κB) and Signal Transducers and Activators of Transcription (STATs)^4, 6, 7^ following an inflammatory signal. NF-κB p65 is a member of the signal-activated transcription factor NF-κB, which forms homo- or heterodimers.^8^ Specifically, p65 forms a heterodimer with another member, NFKB1 (p50), where it serves as the master regulator of proinflammatory response,^9, 10^ cell survival, and proliferation in different cell types, including monocytes, macrophages, dendritic cells, and cancer cells.^10, 11^

Upon macrophage recognition of pathogen-associated molecular patterns (PAMPs) such as the bacterial lipopolysaccharide (LPS), which is a component of the Gram-negative bacterial cell wall,^1, 12^ they become activated. Specifically, Toll-like receptor 4 (TLR4) on these cells dimerizes,^1^ resulting in canonical signaling through the myeloid differentiation factor 88 (MyD88) adaptor protein, which culminates in activation of IκB kinases (IKKα and IKKβ), their phosphorylation of inhibitor of kappa B (IκB), and its degradation. This results in the nuclear translocation of p65 and its transcriptional upregulation of proinflammatory genes, including IL-6, IL-12, and TNF-α, among others,^13–16^ to amplify the inflammatory response.

LPS also activate the antiviral pathway following its endocytosis, resulting in IRF3 phosphorylation, nuclear translocation, and production of type-1 interferons (IFN-I) that activate STAT1.^17^ Specifically, IFN-I binds cognate receptors on cells to activate Janus kinases (JAKs), which phosphorylate STAT1 at Y701. STAT1 then forms homodimers or heterodimerizes with STAT2, and translocates to the nucleus, where it drives the expression of interferon-stimulated genes (ISGs).^18, 19^

Beyond activating transcription factors, pathogenic or inflammatory stimuli are also known to drive the metabolic reprogramming of activated macrophages and other immune cells, which support their inflammatory and bactericidal activity. ^20–23^ In particular, glycolytic reprogramming has been tightly linked to immune and inflammatory activation in response to PAMPs.^24–26^ Macrophages in response to LPS, exhibit a switch from mitochondrial respiration to aerobic glycolysis or the Warburg effect,^24^ such that under aerobic (oxygen-rich) conditions, glycolysis remains the preferred energy source and not oxidative phosphorylation to support their inflammatory functions.^25, 26^ Interestingly, this LPS-induced metabolic switch is also present in M1 macrophages, which are involved in inflammatory responses at inflamed sites and in several inflammatory diseases. Therefore, LPS stimulation of macrophages is a tool to understand both the response of these immune cells to pathogenic Gram-negative bacterial infection and M1 macrophage activity during inflammation.

While several studies have begun to elucidate how LPS-induced metabolic alterations are coupled to post-translational modifications of histones at several inflammatory genes to facilitate their expression,^27–30^ little is known about the precise role of increased glycolysis in coordinating and facilitating p65 and STAT1 activation, which are major transcription factors driving the expression of these inflammatory genes and ISG expression in inflammatory macrophages. There is therefore a need to understand how glycolysis regulates p65 and STAT1 during macrophage activation by LPS, especially given that both p65 and STAT1 co-regulate the expression of several genes, including TNF-α,^31^ nitric oxide synthase (Nos2) in macrophages and ISGs such as CXCL10 in neutrophils.^32, 33^

The events that culminate in the early activation and nuclear translocation of p65 have been extensively studied and well defined during TLR4 signaling by LPS. Interestingly, however, LPS was shown to also promote the expression of p65, which was required for enhanced or prolonged nuclear occupancy of p65 and overall NF-κB signaling in murine macrophages, to enable them to mount an effective immune response against pathogens.^34^

Despite this, the expression of p65 and its kinetics have not been defined in human macrophages, nor is it known whether and how metabolic reprogramming in macrophages regulates p65 expression in response to LPS. This is especially important question given that glycolysis is enhanced under hypoxia, but hypoxia has not been modeled in several in vitro studies despite the fact that many sites of infection, such as the gut, skin epithelial, kidney medulla, vagina mucosa, among others, which also harbor macrophages (resident or monocyte-derived), exist in a physiological hypoxia state (1-2% O_2_), ^35–42^ including several inflammatory sites which are also hypoxic.^43–48^

In this study, we found that LPS stimulation increased the expression of a key glycolytic enzyme, PFKFB3, which we demonstrated is required for late p65 protein expression and phosphorylation, as well as for STAT1 activation in human MDMs. Using a combination of molecular, genetic, and pharmacological approaches, we showed that LPS-induced p65 and select gene-set protein expression rely on the time-dependent upregulation of PFKFB3 under physiological hypoxia and identified the partial involvement of the histone acetyltransferase p300 as a mediator of p65 expression under this condition. We also identified that multiple LPS-activated factors, including STAT1/NF-κB/IRF5, temporally regulate PFKFB3 expression. Overall, we uncovered a positive feedback loop in which p65 and STAT1 activation are driven or sustained by PFKFB3-driven glycolysis in macrophages responding to inflammatory stimuli, which we show feeds back to regulate these factors.

## RESULTS

### LPS Induces Delayed NF-κB p65 Protein Expression in human MDMs

In mouse macrophages, LPS signaling was shown to drive an increase in p65 expression, which was necessary for an effective immune response by increasing NF-κB signaling. We therefore asked if p65 was LPS-inducible in primary human monocyte-derived macrophages (MDMs) under both physiological hypoxia and normoxia. Our results showed that LPS significantly increased p65 protein expression under both hypoxia and normoxia. This increase in p65 protein was accompanied by an increase in its phosphorylation at Serine 536 (S536) (**Figure 1a, b)**, which is a modification associated with increased p65 transcriptional activity.^49^ Corresponding to increased p65 protein expression and phosphorylation, we observed concomitant protein expression of p65-target genes including CD38, CD40, ICAM, and CD44 (**Figure 1c, d)**. This target proteins, including CD38 and CD40, were impaired in cells treated with BMS345541 (BMS), a selective inhibitor of the IKK complex that promotes NF-κB p65 activation and nuclear translocation, and therefore underscores the role of p65 in the induction of this targets **(Figure S1a)**. Next, we assessed the temporal kinetics of p65 expression by performing a time-course experiment in primary human MDMs. Cells were treated with LPS for 1h, 3h, 6h, and 16h, respectively, and p65 expression, phosphorylation, and target gene expression were assessed by western blot. Our results clearly demonstrate that p65 protein expression is only significantly increased after 6h, specifically at the 16h timepoint (**Figure 1e, f)**, albeit we did see increased p65 levels at 12h post-LPS stimulation (data not shown), indicating a late or delayed expression. This observation contrasts with the reported kinetics of p65 protein expression in primary mouse BMDMs, where LPS induces p65 protein expression from 4h, and from 8h in macrophage cell lines.^34^ Additionally, we observed robust protein expression of p65-target genes, CD38 and CD40, at 16h. In contrast, the expression of CD142/tissue factor, another p65 target gene, was restricted to 3h-6h post-LPS stimulation, and wanes by 16h (**Figure 1g)**. These findings reveal distinct dynamics of p65, where its initial early activation within 15 minutes **(Figure S1b)** explains the observed increase in CD142 protein from 3h (early gene), while increased p65 protein expression and phosphorylation at later times (12h-16h) corresponded to expression of late-induced p65 target proteins (**Figure 1g)**. This conclusion is supported by previous work showing that while p65 rapidly translocates into the nucleus upon stimulation, it initially binds promoter regions of certain target genes, while binding to other gene promoter regions occurs later on when they become accessible.^30^ Next, to better understand the dynamics of p65 protein expression, we asked if RELA (p65) mRNA induction precedes the increase in its protein expression. To answer this, we performed qRT-PCR on cells treated with LPS for 3h, 6h, 9h, and 12h. We chose these time points to capture transcriptional changes that would precede translation of the protein of interest. Our results reveal that RELA mRNA gradually increases from 3h and is significantly upregulated at 9h and 12h (**Figure 1h)**, and therefore precedes the increase in p65 protein at 16h. To further demonstrate that new protein synthesis is necessary for p65 protein expression, we inhibited protein translation with the translation inhibitor cycloheximide (CHX). Cells were mock- or LPS-treated for 16h, during which CHX was added at 6h, 8h, or 10h during the course of LPS stimulation. Our results indicate that inhibiting protein translation significantly diminished p65 protein expression **(Figure S1c, d)**. This finding demonstrates that new protein synthesis is required for maximal increase in p65 protein expression, as increased RELA mRNA accumulation fuels the increase in p65 protein expression. Collectively, our findings show that p65 is upregulated at the mRNA and protein level in LPS-stimulated human MDMs under hypoxia and normoxia.

**Figure 1.**
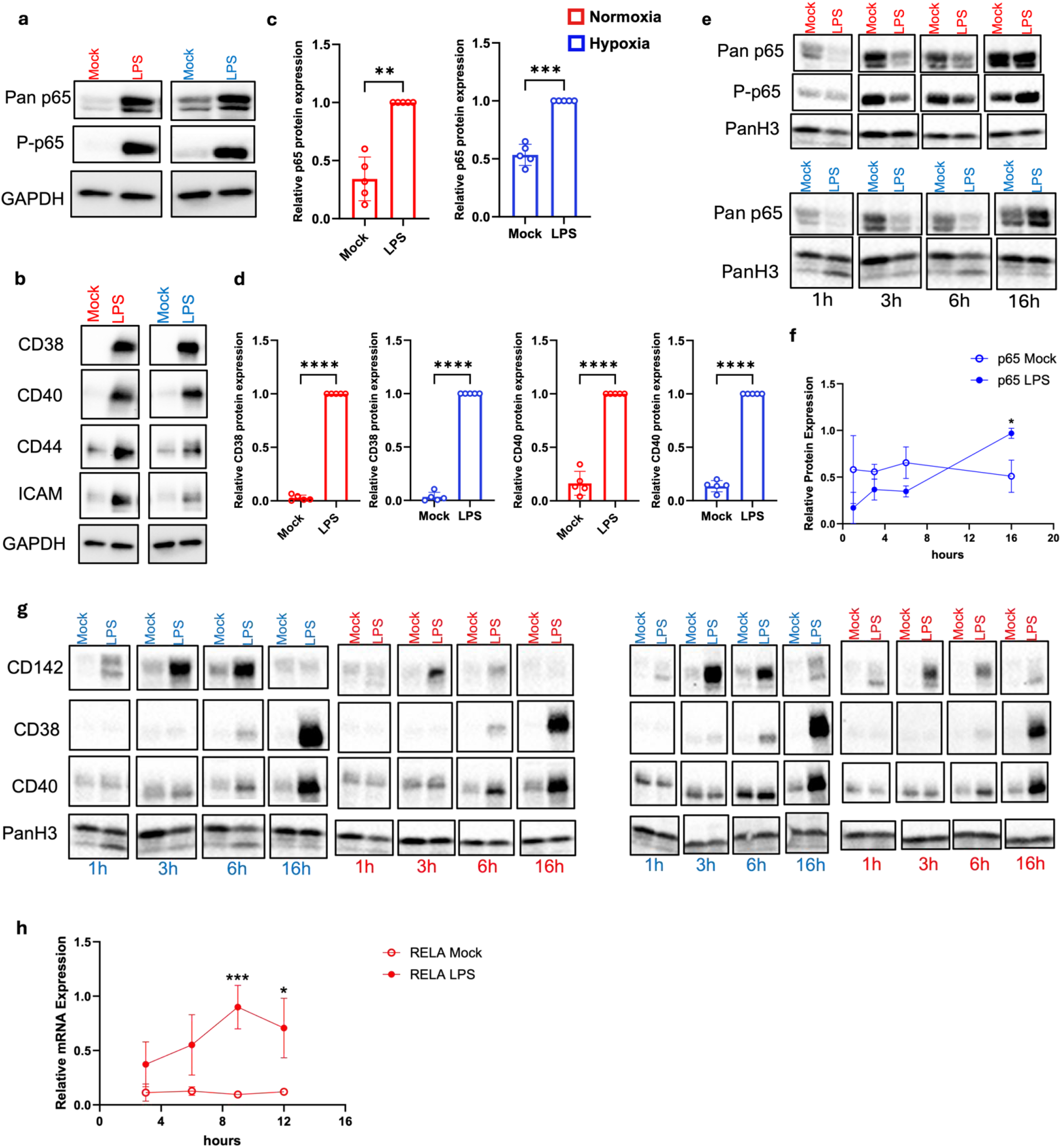
LPS Induces Delayed p65 Protein Expression in human MDMs. Human monocyte-derived macrophages (MDMs) were left untreated (mock) or stimulated with LPS (10ng/ml) for 16h under normoxia (red) and hypoxia (blue). Immunoblot was performed on total cell lysate, with a representative blot shown for (a) Total p65 (Pan p65) and phosphorylated p65 at S536 (P-p65), and (b) p65 target genes indicated: CD38, CD40, CD44, and ICAM, using their specific antibodies. GAPDH served as a loading control. Densitometric analysis is shown in 5 independent donors for (c) Pan p65 and (d) CD38 and CD40 expression in Mock versus LPS-stimulated cells under normoxia and hypoxia. (e) Human MDMs were mock or stimulated with LPS for the indicated time points under normoxia (red) and hypoxia (blue). Immunoblot was performed with representative blot shown for Pan p65, P-p65 and (g) CD142, CD38, and CD40 protein expression kinetics with 2 representative donors shown. (f) Densitometric analysis of 3 independent donors for p65 protein kinetics in (e) is shown for hypoxia. PanH3 was used a loading control. (h) qRT-PCR was performed on Mock versus LPS treated human MDMs over 3h, 6h, 9h, and 12h. Graph represents Relative p65 mRNA expression in 4 independent donors normalized to GAPDH. Statistical significance was determined using Student’s paired T-test; *p<0.05, **p<0.01, ***p<0.001, ****p<0.0001. Error bars indicate SDs.

### LPS Drives a Time-Dependent Induction of Key Glycolytic Enzyme, PFKFB3

Next, we sought to understand the mechanism by which LPS drives a delayed increase in p65 protein expression in human MDMs. Stimulus-inducible inflammatory genes are classified into two groups based on their independence (early genes) or dependence (late genes) on chromatin remodeling.^9, 30, 50^ Early genes are also marked by CpG (cytosine-phosphoguanosine) patches^50, 51^ and, since they do not require remodeling, are readily accessible for immediate transcription in response to inflammatory stimuli such as LPS.^30^ In contrast, late genes are marked by low CpG content,^50, 51^ absence of active histone marks,^30^ and are often inaccessible for transcription factor binding unless they undergo remodeling.^50, 52^ As a consequence, their transcription is not immediate; thus the term “late genes”.^30^ Interferon regulatory factors (IRFs) are a family of transcription factors essential for chromatin remodeling, acting as pioneer transcription factors or co-activators that recruit both ATP-dependent remodelers such as the SWI/SNF family^53^ and histone modifying enzymes to chromatin.^54^ Since the human REL A/p65 gene has an Interferon regulatory factor-1 (IRF-1) binding site,^55^ we asked if IRF-1 was required for p65 expression. Thus, we performed siRNA knockdown of IRF-1 for 16h and then treated the cells with LPS for another 16h. Our result suggest that IRF-1 expression is dispensable for p65 protein expression (**Figure 2a, b)**. Therefore, other remodeling-dependent mechanisms underlie the LPS-dependent delayed increase in p65 mRNA and protein levels.

**Figure 2.**
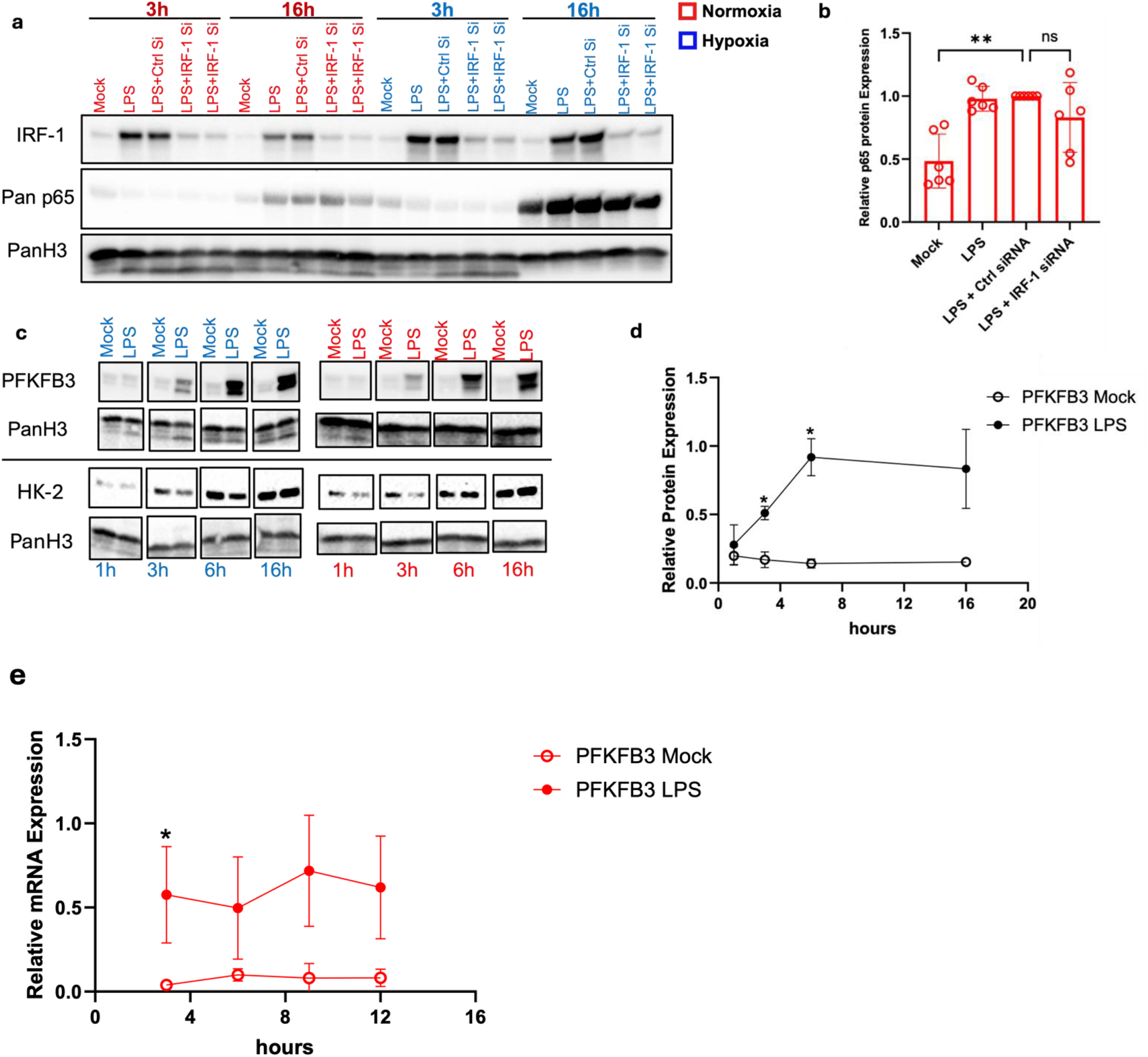
LPS Drives a Time-Dependent Induction of Key Glycolytic Enzyme, PFKFB3. Human MDMs were transfected with control siRNA (Ctrl Si) or IRF-1 siRNA for 16h under normoxia. Thereafter, cells were stimulated with LPS for either 3h or 16h under normoxia (red) and hypoxia (blue), respectively. (a) Immunoblot of IRF-1 and Pan p65 is shown and PanH3 served as a loading control. (b) Relative p65 expression under normoxia in 6 independent donors is quantified by densitometry. (c) Representative Immunoblot of PFKFB3 and Hexokinase 2 (HK-2) from the time-course assay in Mock versus LPS-stimulated cells for the indicated time points is shown for normoxia (red) and hypoxia (blue). (d) Densitometric analysis in 3 independent donors for PFKFB3 protein kinetics under hypoxia is shown. (e) qRT-PCR was performed on Mock versus LPS-treated human MDMs over 3h, 6h, 9h, and 12h under normoxia. Graph represents Relative PFKFB3 mRNA expression in 4 independent donors normalized to GAPDH.

LPS exposure triggers the metabolic rewiring of mouse BMDMs towards glycolysis and promotes histone acetylation at specific inflammatory genes.^29, 56^ Consistently, in response to external stimuli, gradual increases in histone methylation and acetylation activation marks such as H3K4 trimethylation (H3K4me3) and H3K27 acetylation (H3K27Ac) have been reported at the chromatin of late genes.^30, 57, 58^ These modifications are often fueled by nutrient-derived metabolites produced during the metabolic reprogramming of cells,^59^ such as macrophages responding to a pathogen or their product. This LPS-induced metabolic rewiring in mouse BMDMs was first characterized by a rapid increase in glycolysis and TCA cycle metabolites,^29^ followed by a commitment to aerobic glycolysis.^25, 26^ Based on this existing body of literature, we therefore assessed the protein expression of key glycolytic enzymes, hexokinase-2 (HK-2), the first rate-limiting enzyme of glycolysis, and 6-phosphofructo-2-kinase (PFKFB3), a key glycolytic enzyme that is necessary for allosteric activation of the second rate-limiting enzyme of glycolysis, 6-phosphofructokinase-1 (PFK-1), and increased glycolytic flux. Our results demonstrated a striking time-dependent increase in the protein level of PFKFB3 in contrast to minimal changes in HK-2 expression in LPS-stimulated cells (**Figure 2c, d)**, suggestive of PFKFB3’s known role in increasing glycolytic flux. We also observed a consistent time-dependent upregulation of PFKFB3 transcripts, with a significant increase at 3h, followed by continued increases at later time points(**Figure 2e)**. Overlaying PFKFB3 protein expression kinetics with the p65 protein expression kinetics reveals that PFKFB3 protein induction precedes the significant increase in p65 mRNA and protein expression **(Figure S2).** Taken together, we demonstrate that LPS stimulation drives a temporal increase in PFKFB3 but not HK-2 expression prior to maximal p65 expression, suggesting that cells may couple changes in cellular metabolism to p65 expression.

### Glycolysis is Required for LPS-Induced p65 Protein Expression

Given our finding that PFKFB3 expression is significantly upregulated at the mRNA and protein levels before maximal p65 induction, we hypothesized that glycolysis is a critical regulator modulating the switch to p65 transcription and induction in LPS-stimulated macrophages. To test this, we employed two mechanistically distinct approaches that limit glycolysis or glucose metabolism in cells. We pretreated cells for 2h with oxamate (LDHA inhibitor) or treated cells in galactose-containing media (an alternative carbon source to glucose), then stimulated cells with LPS for 16h. In agreement with our hypothesis, p65 expression, phosphorylation, and other late proteins (CD38, CD40) were all significantly reduced in both oxamate-(**Figure 3a, b)** and galactose-cultured LPS-stimulated cells only under hypoxia (**Figure 3c, d, S3a)**. In contrast, normoxia masks these effects, as p65 expression was not impaired by inhibiting glycolysis (**Figure 3c, d, S3a)**. Next, given that PFKFB3 was significantly increased after LPS stimulation, we reasoned that inhibiting glycolysis after LPS treatment or within the early time-frame of PFKFB3 expression (1h-6h) would still inhibit p65 protein expression. To do this, we pre-treated cells with oxamate for 2h (t = −2h) (standard) or added oxamate either 1h, 3h, or 6h post-LPS stimulation of cells. Strikingly, our western blot results clearly showed that p65 protein levels were diminished with oxamate at all time points, regardless of when oxamate was added to cells, and this effect was more pronounced under hypoxia as expected **(Figure S3b)**. Our findings thus underscore the critical role of glucose-fueled metabolism, which is an adaptation under hypoxia, in mediating p65 protein and target gene expression in response to LPS. Next, we wanted to determine whether PFKFB3 is directly necessary for p65 expression. To do this, human MDMs were pretreated with the PFKFB3 inhibitor, Pfk15, for 2h prior to stimulation with LPS for 16h. Indeed, Pfk15 significantly reduced the expression of p65, CD38, and CD40 under hypoxia. However, only ∼55.5% of subjects showed a slight reduction in p65 protein levels under normoxia (**Figure 3e, f)**. Consistently, p65 mRNA expression was not impaired in Pfk15-treated cells under normoxia (**Figure 3g)**. To further confirm the activity of the Pfk15 compound, we performed siRNA knockdown of PFKFB3 in human MDMs for 16h. PFKFB3 knockdown was followed by LPS stimulation for another 6h and 16h, respectively. We confirmed knockdown of PFKFB3 at both 6h and 16h by western blot, and this corresponded to a modest and consistent decrease in p65 expression and phosphorylation under normoxia and hypoxia at 16h (**Figure 3h, i)**. Overall, these findings support our hypothesis that glycolysis is required for p65 expression, especially under hypoxia, and uncovered PFKFB3 as a key glycolytic regulator of p65 expression in human MDM activated with LPS.

**Figure 3.**
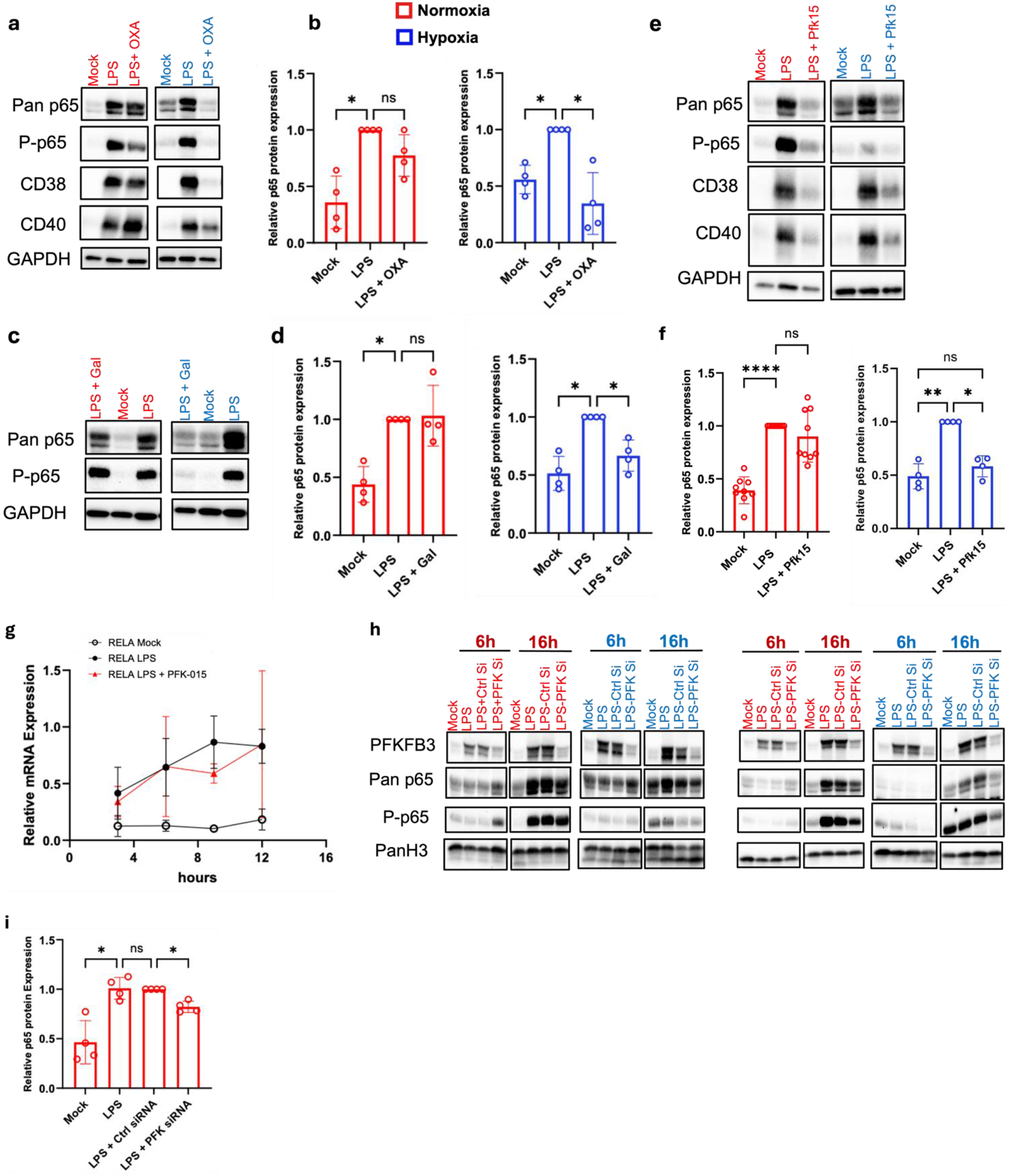
Glucose Metabolism is Required for p65 and late genes expression in LPS-Stimulated Human MDMs. Human MDMs were pre-treated with a glycolytic inhibitor (a) 10 mM oxamate (OXA) for 2h, and (c) alternatively, MDMs were cultured in a no-glucose medium supplemented with 11.1 mM galactose (Gal). (a, c) Cells were then stimulated with LPS for 16h, and an Immunoblot was performed to detect Pan p65, P-p65, CD38, and CD40. GAPDH served as a loading control. (b, d) Densitometric analysis of Pan p65 from 4 independent donors in a and c experiments is shown. (e) Relative p65 mRNA expression was quantified in human MDMs pre-treated with PFK15 for 2 h and stimulated with LPS for 3h, 6h, 9h, and 12 h, respectively. Statistical significance was determined using Student paired T-test; *p<0.05, **p<0.01, ***p<0.005. (f, g) Human MDMs were mock- or pre-treated with PFKFB3 inhibitor (PFK15 – 5 μM) for 2h, then stimulated with LPS for 16h. (f) A representative Immunoblot for the detection of Pan p65, P-p65, and CD38 is shown. GAPDH served as a loading control, and (g) Densitometric analysis for Pan p65 expression is depicted under normoxia (red) and hypoxia (blue). (h) Human MDMs were transfected with control siRNA or PFKFB3 (PFK) siRNA for 16h under normoxia and then mock- or stimulated with LPS for either 6h or 16h under normoxia (red) and hypoxia (blue), respectively. Immunoblots of 2 independent donors (A and B) is shown for detection of PFKFB3, Pan p65 and P-p65, while PanH3 served as loading control. (i) Densitometric analysis of Panp65 expression under normoxia is shown. Statistical significance was determined using repeated-measures One-Way ANOVA, and post hoc with Šídák’s multiple comparisons test; *p<0.05, **p<0.01, ***p<0.001, ****p<0.0001. Error bars indicate SDs.

### Downstream Glycolytic Metabolite, Pyruvate, is a Player in LPS-Induced p65 Expression

To further understand and investigate the regulation of p65 expression by glycolysis, we performed addback experiments with the glycolytic metabolite, pyruvate, in oxamate-treated cells. Oxamate inhibits LDHA activity by outcompeting pyruvate, a downstream glycolytic metabolite, thereby preventing lactate production and the recycling of NAD+ (**Figure 4a)**, ultimately impairing the glycolytic flux. Knowing this, we hypothesized that adding pyruvate exogenously would restore p65 expression under hypoxia where glycolysis was required. To this end, we performed addback experiments with exogenous pyruvate. Briefly, cells were pre-treated with oxamate for 2h, then treated concomitantly with LPS and exogenous pyruvate (Exo-Pyr) for 16h. Consistent with our hypothesis, oxamate significantly reduced p65 protein expression, and Exo-Pyr rescued this under hypoxia. In addition, p65 target genes such as CD38 were also slightly rescued, as shown by western blot (**Figure 4b, c, S4a)**. Altogether, our results demonstrate that the glucose-derived metabolite, pyruvate, can act downstream of glycolysis to facilitate LPS-induced expression of p65.

**Figure 4.**
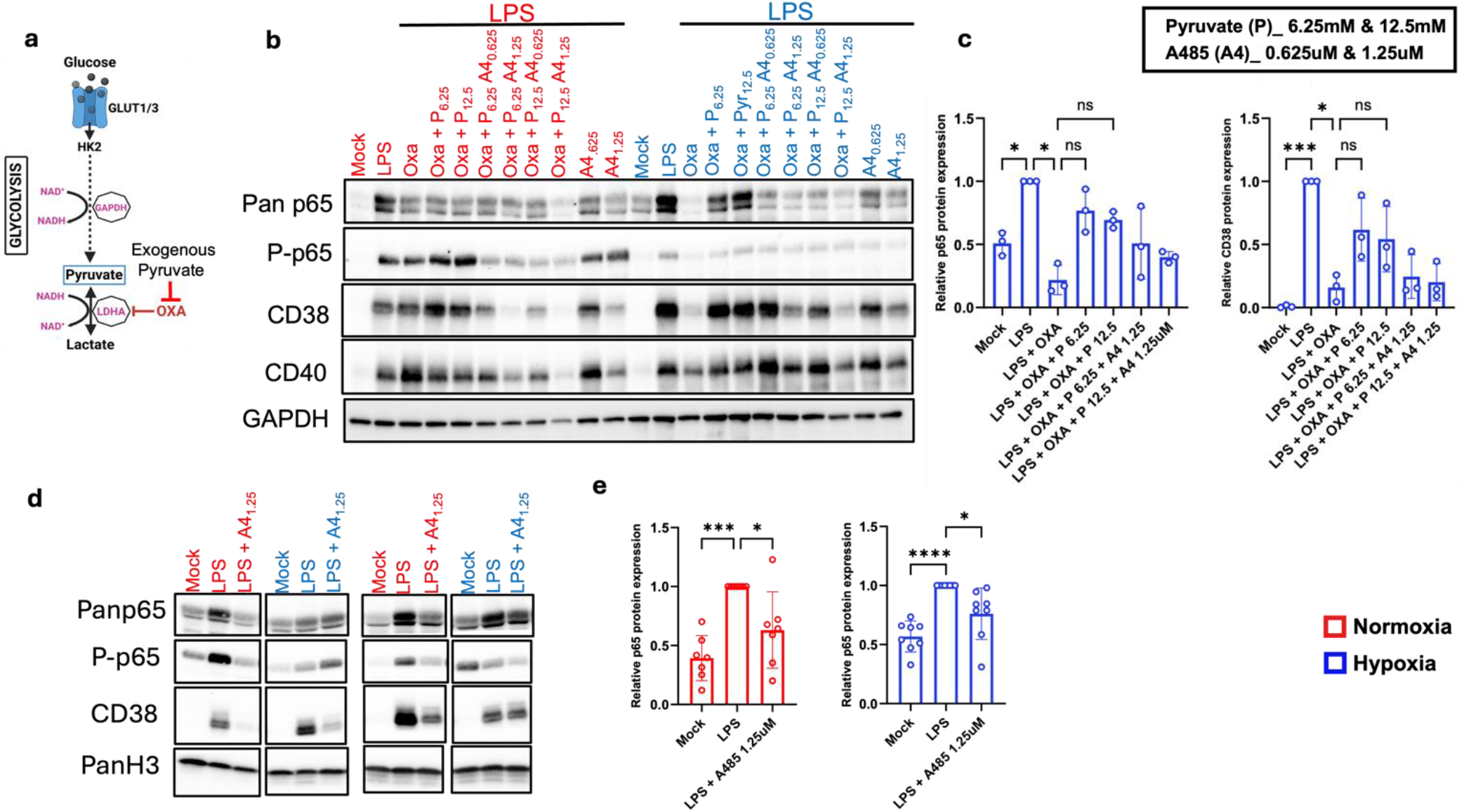
p300 Activity Partly Mediates p65. (a) Human MDMs were mock- or pre-treated with 10 mM oxamate (OXA) or p300 inhibitor A485 (shortened as A4), respectively, at the indicated concentration for 2h. Afterwards, LPS and exogenous pyruvate (Pyr) at 6.25 mM and 12.5 mM, respectively, were added to cells for 16 h. Immunoblotting was performed to detect Pan p65, P-p65, CD38, and CD40. GAPDH served as a loading control. (b) Densitometric analysis for relative Pan p65 and CD38 expression under hypoxia is shown. Human MDMs pre-treated with 1.25 μM A4 for 2h and stimulated with LPS for 16h under normoxia and hypoxia were subjected to immunoblot assay for Pan p65, P-p65 and CD38. (c) Data from 2 representative donors are shown, and (d) Densitometric analysis for relative Pan p65 expression is shown. Statistical significance was determined using repeated-measures one-way ANOVA and post hoc with Šídák’s multiple comparisons test; *p<0.05, **p<0.01, ***p<0.001, ****p<0.0001. Error bars indicate SDs.

### p300 Activity is Partly Required for p65 Expression in LPS-Stimulated Macrophages

We then asked the next natural question. What is the mechanism by which glycolysis, or its metabolite, drives p65 expression in LPS-stimulated human MDMs? Increases in histone acetylation and lactylation license genes to be transcriptionally ready, and histone acetylation has been shown to increase on late-expressed genes in response to inflammatory stimuli.^30^ Additionally, a PFKFB3-dependent increase in lactate and histone lactylation on genes such as RELA (p65) has been reported in chronic kidney disease,^60^ which is characterized by low-grade inflammation.^61, 62^ Interestingly, our results also showed that inhibiting glycolysis with oxamate reduced global histone PTMs associated with gene transcription,^63–67^ specifically H3K9 lactylation (H3K9la), H3K4me3, and H3K27ac, in LPS-stimulated human MDMs, which were rescued by exogenous pyruvate **(Figure S4a, b, c, 4b)**. The restoration of these PTMs by Exo-Pyr correlates with p65 protein rescue by Exo-Pyr in oxamate-treated cells (**Figure 4b, c, S4a)**. These findings suggest that glycolysis-fueled histone post-translational modification may underlie LPS-induced p65 expression in human MDMs, but this needed to be tested.

Histone PTMs are catalyzed by histone-modifying enzymes, which covalently conjugate metabolite-derived modifications to histone and non-histone proteins.^59^ Since the histone acetyltransferase (HAT) p300, is a known writer for the H3K27ac mark^68^ and has been recently shown to mediate histone lactylation, reviewed in,^69^ including H3K9la in microglia and neural stem cells,^64, 70^ we asked if p300 acts downstream of glycolysis and its metabolites to facilitate p65 expression. To determine this, cells were pre-treated with A485 (a canonical inhibitor of p300 catalytic activity) and then stimulated with LPS for 16h. Our results showed that A485 only partially diminished LPS-induced p65 expression in human MDMs under hypoxia, with more pronounced diminution observed under normoxia (**Figure 4d, e)**. In addition, p65 target genes, CD38 and, to a lesser extent, CD40 (late genes induced alongside p65), were also slightly impaired under hypoxia. Consistent with our findings that p300 is necessary for p65 expression, A485 reduced Exo-Pyr-dependent rescue of p65 and CD38, albeit not statistically significant (**Figure 4b, S4a)**. In sum, our findings suggest that LPS stimulation couples increases in glycolysis and its metabolites to p65 expression, partly through the activity of p300.

### LPS-Induced TBK1/IKKƐ Kinase Activity Promotes Early and Sustained STAT1 S727 Phosphorylation

In addition to LPS activating the IKK signaling that culminates in early NF-κB p65 activation and nuclear translocation, LPS also activates the STAT1 antiviral pathway.^71^ This can be directly through S727 phosphorylation^31, 71^ and indirectly through autocrine/paracrine type-1 IFN signaling (IFN-I), which drives STAT Y701 phosphorylation by activating JAKs.^18, 19^ Given the extensive evidence of crosstalk between the NF-κB and STAT1 pathways, including co-regulation of some inflammatory genes and some ISGs,^72–75^ we sought to determine whether this crosstalk extended to the metabolic regulation of STAT1. To answer this question, we first determined the kinetics of STAT1 in primary human MDMs. Consistent with previous reports in mouse BMDMs,^31^ LPS caused rapid phosphorylation of STAT1 at its transactivation domain (S727) within 15 min in primary human MDMs (**Figure 5a)**. Through a time course experiment, we also determined that STAT1 becomes phosphorylated at its Y701 residue within an hour, a modification required for STAT1 homo- or heterodimerization and subsequent nuclear translocation. This was accompanied by increased S727 phosphorylation at 1h (**Figure 5b)**. Interestingly, the kinetics of STAT1 S727 and Y701 phosphorylation were slightly delayed in human PMA-differentiated THP-1 macrophages, occurring at 1h and 3h, respectively **(Figure S5a)**. Interestingly, BX795, the dual kinase inhibitor for TBK1/ IKKƐ kinase, which was previously shown to inhibit IFN-I production by blocking IRF3 phosphorylation and nuclear translocation,^76^ also inhibited STAT1 S727 phosphorylation and, expectedly, Y701 phosphorylation given its dependence on IFN-I signaling, across all time points examined under normoxia and hypoxia (**Figure 5b)**. As expected, BX795 also reduced the expression of STAT1-dependent genes/ISGs such as IRF-1, MDA5, and CXCL10, as well as CD38 and CD40 (**Figure 5b, S5b)**, some of which are known to be co-regulated by NF-κB. Altogether, these findings demonstrate that STAT1 S727 and Y701 phosphorylation are induced early with distinct kinetics and are regulated by the TBK1/ IKKƐ kinases throughout the course of LPS exposure.

**Figure 5.**
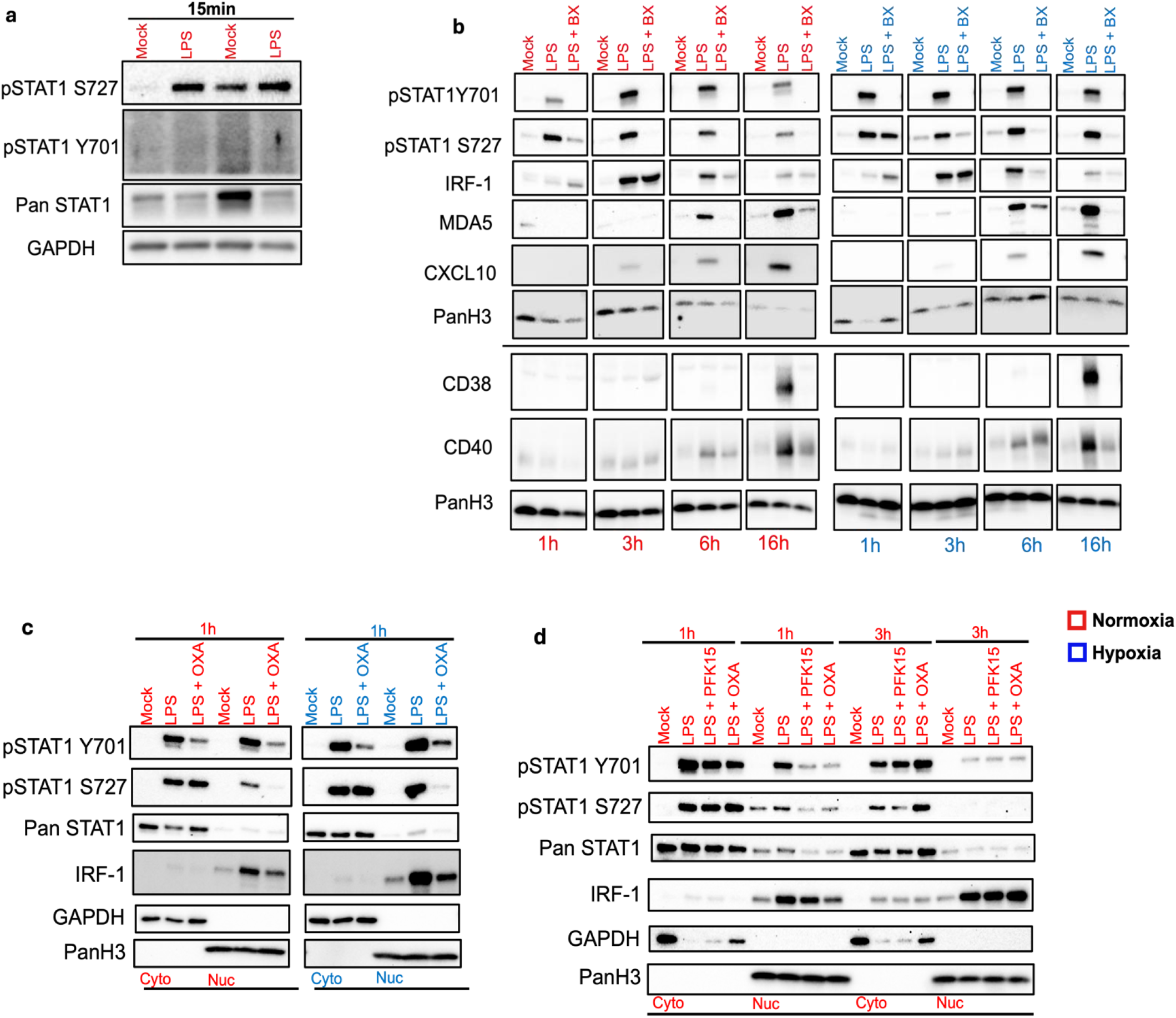
Glucose Metabolism is Required for Early STAT-1 Y701 Phosphorylation and IRF-1 Expression. **(a)** Human MDMs were mock- or LPS-stimulated for 15 min. Immunoblot showing STAT1 phosphorylation at S727 and Y701 in 2 independent donors under normoxia. **(b)** Human MDMs were pre-treated with TBK1/IKKƐ inhibitor, BX795 (BX), for 2h, thereafter stimulated with LPS for the indicated time points. Immunoblot analysis of STAT-1 phosphorylation and known target genes is shown. Data are representative of 2 donors. **(c)** Primary Human MDMs were pre-treated with oxamate (OXA) for 2h, thereafter stimulated with LPS for 1h under normoxia (red) and hypoxia (blue). Immunoblot analysis of nuclear fractionated cells showing STAT-1 phosphorylation at Y701, S727, Pan STAT1, and IRF-1. GAPDH and PanH3 were used as controls for cytoplasmic and nuclear fractions, respectively. **(d)** Immunoblot analysis of nuclear fractionated cells pretreated with OXA or PFK15 for 2h and stimulated with LPS for 1h and 3h respectively, under normoxia. STAT-1 phosphorylation at Y701 and S727, Pan STAT-1, and IRF-1 were probed using their respective antibodies. GAPDH and PanH3 were used as controls for cytoplasmic and nuclear fractions, respectively.

### Glucose Metabolism is Required for the LPS-Induced Rapid Activation and Nuclear Translocation of STAT1

Next, we tested the role of glucose metabolism in STAT1 activation and nuclear translocation. To do this, we pretreated human MDMs with the glycolytic inhibitors oxamate or PFK15 for 2h, then stimulated them with LPS for 1h. Our nuclear fractionation results indicate that both inhibitors significantly diminished STAT1 phosphorylation at Y701, but only a modest diminution was observed at S727 (**Figure 5c**). We also observed that STAT1 nuclear translocation was significantly diminished irrespective of STAT1 S727 phosphorylation status in Pfk15- or oxamate-treated cells, and this can be explained by the loss of Y701 phosphorylation in these cells (**Figure 5c**). These effects were also recapitulated in PMA-differentiated THP-1 macrophages, albeit with slightly different kinetics (**Figure S5c, d**). In addition to STAT1 activation, we asked whether the STAT1 target gene, IRF-1, is also regulated by glucose metabolism. We stimulated human MDMs pretreated with PFK15 or oxamate with LPS for 1h and 3h, respectively, to monitor STAT1 and IRF-1 levels in the nucleus. Consistent with oxamate and PFK15 treatment causing reduced levels of nuclear-translocated STAT1 at 1h, the protein level of IRF-1 in the nuclear fraction was equally diminished under normoxia at 1h but not 3h, whereas IRF-1 levels remained diminished for up to 3h under hypoxia in the nuclear fraction (**Figure 5d, S5e, f**). Taken together, our findings demonstrate that early STAT1 activation and nuclear translocation, coupled with IRF-1 expression, are glucose-dependent steps. To our knowledge, this is the first time that glucose metabolism has been directly shown to be important for STAT1 phosphorylation and nuclear import in macrophages. Previous studies have mostly demonstrated that high-glucose conditions enhance STAT1 phosphorylation and transcriptional activity under different conditions and in cells, including macrophages, cardiac fibroblasts, and glomeruli of diabetic mice, among others.^77–79^ Altogether, our findings demonstrate that PFKFB3-driven glycolysis is also functionally coupled to the canonical STAT1 pathway activation and downstream responses during TLR4 signaling by promoting STAT1 Y701 phosphorylation and nuclear translocation.

### LPS-Dependent PFKFB3 Expression is Temporally Regulated by Multiple Factors

Our findings have uncovered how glucose metabolism driven by enhanced PFKFB3 expression coordinates p65 expression and STAT1 activation during innate immune signaling by LPS. Thus, we asked which upstream pathway, or factor, regulates the time-dependent induction of PFKFB3 in LPS-stimulated human MDMs. LPS activates several pathways, both directly and indirectly, in activated cells. TBK1/IKKƐ signaling, for example, was recently shown to promote rapid metabolic reprogramming in BMDMs^56^ and previously, in dendritic cells responding to LPS and other PAMPs.^80^ But their role in driving glycolytic gene expression, such as PFKFB3, has not been examined. LPS also rapidly activates p65 through the activity of the kinases IKKα & IKKβ. In the context of PFKFB3 expression, p65 has been shown to promote HIF-1α-dependent upregulation of PFKFB3. Additionally, TNFα-induced p65 phosphorylation was associated with increased PFKFB3 expression in endothelial cells,^81^ and p65 was identified as a transcription factor for PFKFB3 in trophoblast co-transfected with LPS.^82^ Also, evidence in the literature has shown that Interferon Regulatory Factor 5 (IRF5), a transcription factor that is activated and imported to the nucleus upon TLR4 signaling,^83^ promotes the expression of certain glycolytic genes, either directly as a transcription factor^84, 85^ or indirectly by activating AKT2.^86^ Based on this body of literature, we queried which of these factors regulates the temporal induction of PFKFB3 in LPS-stimulated human MDMs. To answer this, we pretreated cells with either BMS (IKKα and IKKβ inhibitor), BX795 (TBK1/ IKKƐ/STAT1 inhibitor), or YE6 (IRF5 inhibitor) for 2h, and then stimulated cells with LPS for 3h, 6h, and 16h, respectively, and then performed a western blot. Similarly, we performed qRT-PCR on cells stimulated with LPS for 3h, 6h, 9h and 12h to elucidate the transcriptional regulation of PFKFB3 during BX795 and YE6 inhibition, which are not known regulators of PFKFB3.

Our result demonstrated that while BMS significantly diminished PFKFB3 protein expression at 3h but not 6h and 16h (**Figure 6a, f)**, BX795 significantly reduced PFKFB3 protein expression at 6h and 16h but not at 3h post-LPS stimulation (**Figure 6a, b)**, whereas YE6 treatment only significantly reduced PFKFB3 protein expression 16h, with a slight diminution at 6h (**Figure 6a, d)**. Altogether, this indicates a time-dependent regulation of PFKFB3 protein expression by these proteins. Our qRT-PCR corroborates these findings and demonstrates that while BX795 did not inhibit PFKFB3 expression at 3h post-LPS stimulation, it downregulates PFKFB3 at 6h, 9h, and 12h, with statistical significance at 6h and 12h **(Figure S6a)**. Although YE6 did not affect PFKFB3 mRNA expression at 3h, we observed a modest, non-significant reduction at 9h and 12h post-LPS stimulation **(Figure S6b)**, and this downregulation may underlie the reduction in PFKFB3 we detected by western blot at 16h in Y46-treated cells (**Figure 6c)**.

**Figure 6.**
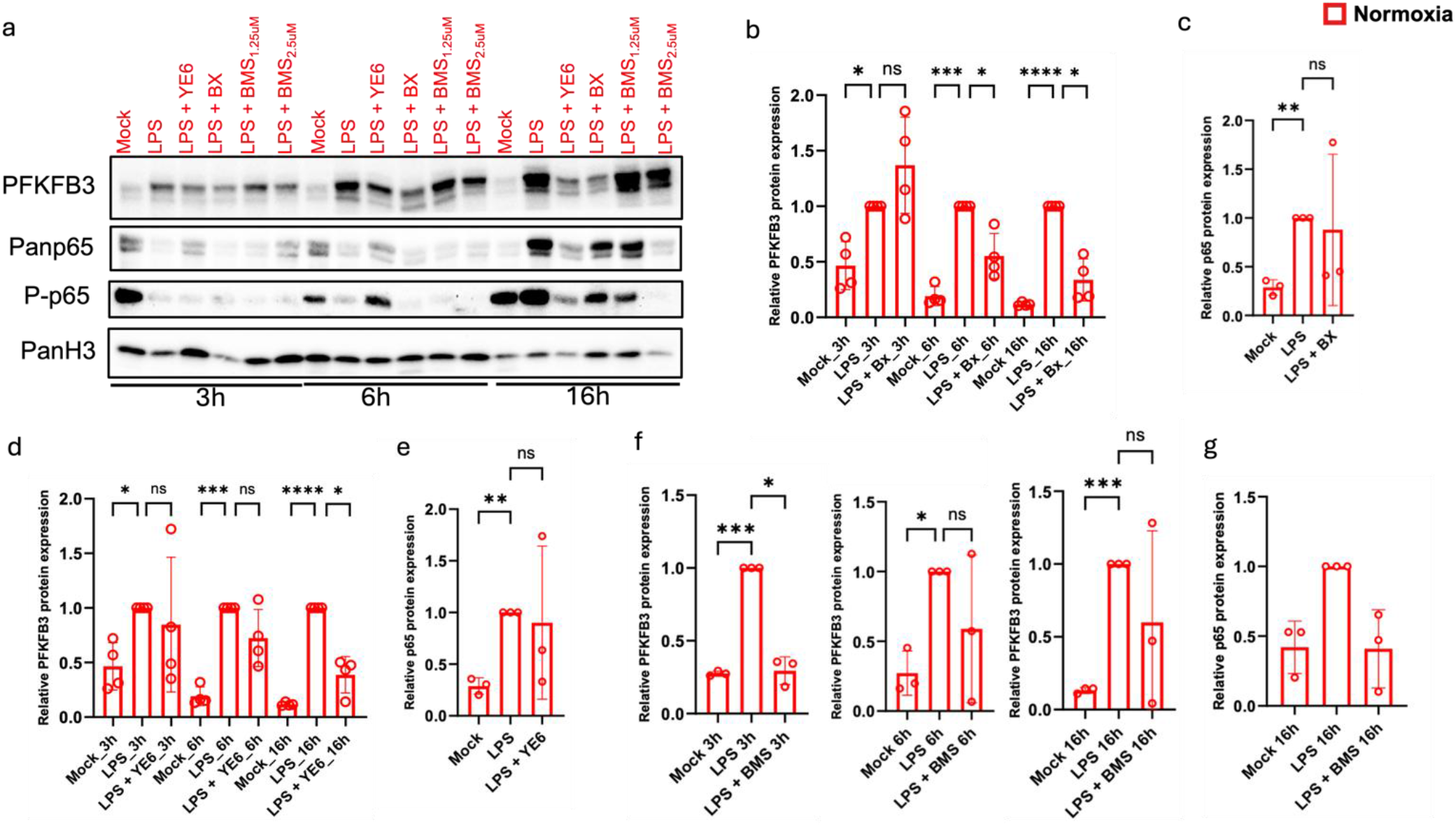
Multiple TLR4-Activated Factors Regulate PFKFB3 Induction in Primary Human MDMs. (a) Representative Immunoblot depicting primary human MDMs pre-treated with BX, YE6, and IKKα and β inhibitors (BMS34451 or BMS - 2.5 μM) and stimulated with LPS for indicated times. (b) Densitometric analysis of PFKFB3 for the indicated times and (c) p65 protein expression for 16h in (a) for BX-treated cells stimulated with LPS. (d) Densitometric analysis of PFKFB3 for the indicated times and (e) p65 protein expression for 16h in (a) for YE6-treated cells stimulated with LPS. (f) Densitometric analysis of PFKFB3 for the indicated times and (g) p65 protein expression for 16h in (a) for BMS-treated cells stimulated with LPS. Statistical significance was determined using repeated-measures One-Way ANOVA, and post hoc with Šídák’s multiple comparisons test*p<0.05, **p<0.01, ***p<0.001, ****p<0.0001. Error bars indicate SDs.

We also assessed the effect of inhibiting these factors on late induction of RELA and demonstrated that RELA/p65 mRNA expression was significantly downregulated at 12 h in BX795- and YE6-treated cells **(Figure S6b, d)**. Surprisingly, we observed that BX795 and YE6 did not always reduce p65 protein expression (**Figure 6c, e)**. Collectively, our findings suggest that LPS-induced PFKFB3 expression is a multifactorial process regulated by the time-dependent involvement of individual LPS-activated factors identified here as TBK1/IKKƐ/STAT1, IKK/p65, and IRF5, which, interestingly, we showed to be regulated by PFKFB3-driven glycolysis (for p65 and STAT1) and therefore suggest a positive feedback loop.

## DISCUSSION

In this study, we modeled hypoxia, which is physiologic for many tissues where gram-negative bacterial infection occurs but also inflammatory sites in different diseases, ^43–48^ to understand the metabolic regulation of inflammatory and antiviral transcription factors during TLR4 signaling. We demonstrate that glucose metabolism is intricately linked to the expression and activation of major transcription factors, NF-κB and STAT1, that drive inflammatory and antiviral responses during LPS stimulation, and promotes macrophage M1 polarization,^87, 88^ which are macrophage subtypes abundant and associated with severity in inflammatory diseases such as rheumatoid arthritis, multiple sclerosis, and inflammatory bowel disease, among others.^89–93^ First, we show that in primary human MDMs, NF-κB p65 is induced and phosphorylated at its S536 transactivation domain in a delayed fashion, and an increase in glycolysis driven by upregulated PFKFB3 is required for this only under hypoxia, not normoxia. In addition, our findings revealed, for the first time, that increased glucose metabolism is necessary for early STAT1 activation, its nuclear translocation, IRF-1 expression, and expression of certain ISGs. Surprisingly, we found that PFKFB3 expression is temporally regulated by a trio of upstream activated pathways, including IKK/p65, TBK1/IKKƐ/STAT1, and IRF5. Our work uncovered that TLR4 signaling, through these factors, triggers positive glycolytic feedback by inducing PFKFB3 and its activity, thereby amplifying the activation of p65 and STAT1 during inflammatory signaling.

The finding that NF-κB p65, encoded by the RELA gene, is inducible by pathogenic LPS concentrations was reported in murine macrophages, where it was demonstrated that it promoted a positive feedback loop that promoted the sustained nuclear occupancy of NF-κB and an appropriate antibacterial response.^34^ Our findings extended this to primary human macrophages and demonstrated that LPS stimulation also upregulated RELA/p65 expression at the transcript and protein levels in a time-dependent manner, but over a timeframe distinct from the reported kinetics of RELA induction in murine macrophages. Specifically, p65 mRNA and protein were strongly upregulated only at 9h and 16h post-LPS stimulation, respectively, in primary human MDMs, whereas at 4h post-LPS stimulation in primary mouse BMDMs.^34^ Correspondingly, we identify a set of gene products known to be co- or independently regulated by NF-κB and STAT1 that were robustly increased alongside p65 at 16h post-LPS stimulation. They include CD38 (an inflammatory marker in M1 macrophages),^94^ CD40 (interacts with CD40L on T-lymphocytes to promote their activation),^95^ ICAM (required for phagocytosis in LPS-stimulated BMDMs and implicated in efferocytosis),^96^ and CD44.^97^ Our finding that p65 target proteins increase alongside p65 expression, and the presence of total and phosphorylated p65 in the nuclear fraction of cells even at 16h post-LPS stimulation, supports the conclusion from previous studies that increased p65 expression increases NF-κB-dependent signaling at the single-cell level in BMDMs.^34^ Indeed, further support comes from the observation that p65 overexpression correlated with increased NF-κB activation and poor patient outcomes in a type of cancer.^98^

Macrophages responding to PAMPs such as LPS undergo cellular metabolic reprogramming, a phenotype tightly linked to their functions, including the production of inflammatory cytokines and microbicidal killing,^20–23^ followed by a transition to an anti-inflammatory state.^20^ The former response is characterized by a rapid glycolytic burst upon stimulation with TLR ligands,^56, 80^ an increase in TCA cycle metabolites,^29^ and finally commitment to a Warburg state termed aerobic glycolysis.^25, 26^ Our study uncovered that the mRNA and protein expression of the second rate-limiting glycolytic enzyme, PFKFB3, were significantly upregulated prior to a significant increase in p65 expression under normoxia and hypoxia. However, strikingly, perturbing glycolysis resulted in a significant reduction in p65 protein expression only under hypoxia, an effect masked by normoxia. This is an interesting observation, despite the known Warburg phenotype reported in mouse BMDMs under normoxia. However, because normoxia (21% O_2_) is also not the physioxia in the blood or tissues in vivo, and given that hypoxia is the more relevant condition to study many inflammatory conditions,^43–48^ the observed glycolytic dependency under hypoxia for expression of p65 and certain downstream target proteins unveils metabolic vulnerabilities that can be targeted to modulate immune response to pathogenic infection or macrophage functions in inflammatory disease. Interestingly, in contrast to PFKFB3, we did not see robust changes in the expression of the first rate-limiting glycolytic enzyme, HK-2, across the time points. This suggests that HK-2 may not be the molecular target of LPS in driving sustained glycolytic reprogramming of cells, even though it has been shown to support a rapid glycolytic burst in dendritic cells by interacting with the mitochondria.^80^ Consistently, increased PFKFB3 expression is one of the four processes necessary for Warburg metabolism in LPS-stimulated macrophages, dendritic cells, and M1 macrophages,^99^ and has also been identified as a driver of the persistent glycolytic phenotype that underlies chronic kidney disease.^60^

NF-κB p65 expression kinetics suggest that it belongs to a class of genes (late genes) that require chromatin remodeling and/or an additional factor before it can be robustly expressed.^50, 52^ Essentially, chromatin remodeling promotes accessibility for transcription factors and co-activators to bind to the promoter of the gene of interest. An important aspect of chromatin remodeling is epigenetic rewiring, involving the post-translational modifications of histones by histone-modifying enzymes, including HATs, which promote acetylation and lactylation at chromatin, a process necessary to license those genes for transcription.^100–104^ These modifications are driven by nutrient-derived metabolites that serve as substrates for histone-modifying enzymes in cells undergoing metabolic reprogramming.^59^ Our finding that PFKFB3 induction, which enhances glycolytic flux, precedes p65 expression, opened an avenue to explore if glucose metabolism facilitates p65 expression through glucose-fueled histone modifications. Indeed, inhibition of glycolysis with oxamate diminished the levels of histone PTMs typically enriched at active genes/enhancers, including H3K4me3, H3K27ac, and H3K9la, which are also involved in gene transcription. We went on to demonstrate that the histone acetyltransferase p300 may partly regulate p65 expression under hypoxia, unlike normoxia, where the effects are more striking. Despite the fact that TLR/TLR4 signaling has been shown to direct p300 to specific genes in a signal-dependent manner,^105^ and is also implicated in the rapid acetylation of various pro-inflammatory genes,^106^ it is possible that p300 collaborates with additional yet-to-be-identified writer(s) in regulating p65 expression under hypoxia. It is also noteworthy that, from our results, increased p65 protein expression corresponded to an increase in p65 S536 phosphorylation at 16h, both in the cytoplasm and nucleus. This modification has been associated with p65’s ability to drive the expression of specific genes.^107, 108^ Interestingly, while IκB negatively regulates p65 by sequestering it in the cytoplasm or even in the nucleus before export, S536 phosphorylated p65 is not inhibited by nuclear IκB.^107^ Taken together, our findings support the premise that increased p65 expression drives increased NF-κB signaling and is regulated by glycolysis under physiological hypoxia.

The STAT1 pathway has been shown to regulate inflammatory response in macrophages^31, 109^ in addition to its canonical role in the ISG response, including co-regulating some of these genes with NF-κB.^72–75^ LPS signaling can directly promote STAT1 phosphorylation at its S727 transactivation domain, independent of autocrine/paracrine type-1 interferon signaling in murine macrophages.^31, 71^ Our initial findings in primary human MDMs and PMA-differentiated THP-1 macrophages were consistent with reports in BMDMs that LPS drives early STAT1 S727 phosphorylation in response to LPS, as revealed by distinct kinetics of S727 (15min) and Y701(1h) phosphorylation in primary human MDMs, but 1h and 3h, respectively, in PMA-differentiated THP-1 macrophages. Surprisingly, glycolysis inhibition preferentially inhibited Y701 phosphorylation and not S727 phosphorylation, resulting in reduced STAT1 nuclear translocation. This was expected as Y701 phosphorylation is required for STAT1 dimerization and nuclear translocation.^110, 111^ However, in contrast to the report that early S727-phosphorylated STAT1 could translocate to the nucleus independent of Y701 phosphorylation,^31^ we did not observe nuclear-translocated STAT1, even though glycolysis inhibition only minimally affected S727 phosphorylation. However, future studies will determine how glucose metabolism preferentially regulates early STAT1 Y701 phosphorylation. We speculate, however, that since STAT1 undergoes various post-translational modifications, such as acetylation, β-hydroxybutyrylation, and pyruvylation, glycolysis inhibition may alter the ratio of PTMs on STAT1, which may affect its ability to be phosphorylated. Importantly, β-hydroxybutyrylation of STAT1, a modification increased by starvation, has been previously shown to inhibit STAT1 phosphorylation and transcriptional activity.^112^ Alternatively, inhibition of glycolysis may prevent the production of type-1 IFNs that activate JAKs, thus preventing STAT1 phosphorylation in LPS-stimulated macrophages. In line with this idea, inhibition of glycolysis has been shown to reduce IFN-1 production following Retinoic Acid-Inducible Gene (RIG-1) activation in monocyte-derived dendritic cells, resulting in increased replication of the different viruses assessed.^113^ Thus, future work is warranted to understand the mechanisms by which glycolysis regulates STAT1 activation, particularly in human MDMs during pathogenic stimulation.

Concomitant with diminished nuclear STAT1, we observed reduced IRF-1 levels in the nuclear fraction of cells treated with glycolytic inhibitors. Our findings demonstrate that IRF-1 rapidly accumulates in the nucleus. However, the absence of IRF-1 in the cytoplasmic fraction and the striking reduction in the nuclear fraction in glycolytic inhibitor-treated cells both suggest that IRF-1 transcriptional or protein expression, and not its nuclear import, is regulated by glucose metabolism. Corroborating our result that STAT1 activation and downstream gene activation are glucose-regulated in human macrophages, Nareika and colleagues also reported that high-glucose media potentiated the JAK-STAT-dependent expression of metalloproteinase 1 in primary and immortalized human MDMs stimulated with IFN-γ and LPS.^78^ We have expanded on this by directly showing that the phosphorylation, activation, and nuclear translocation of STAT1 depend on increased glycolytic flux early in the response to LPS. Surprisingly, we observed that the reduction in IRF-1 expression lasted for 1h under normoxia but was sustained for 3h under hypoxia in primary human MDMs. Though the reason for these differences remains elusive,

Our findings established a central role for increased glucose metabolism in driving p65 expression in hypoxia and STAT1 activation in response to LPS, providing a rationale for investigating the regulation of PFKFB3 expression in LPS-stimulated macrophages. Surprisingly, we identified TBK1/IKKƐ, which regulates STAT1 S727 and Y701 phosphorylation, and IRF5 as factors that temporally regulate PFKFB3 expression alongside IKK/p65, which have been previously linked to PFKFB3 expression. This means that the early (3h), mid (6h), and late (16h) expression of PFKFB3 may be distinctly or cooperatively regulated by these factors. While this conclusion was made based on experiments performed, we suspect that they may be the same or even exaggerated under hypoxia since the temporal kinetics of PFKFB3 induction were the same under both hypoxia and normoxia. Interestingly, IRF5 has been shown to interact with p65 to target certain inflammatory genes with unique noncanonical sequences in LPS-stimulated GMCSF-derived BMDMs.^114^ Whether this applies to metabolic genes like PFKFB3 is not known. Moreover, IKKβ has also been shown to phosphorylate IRF5, activating it.^115, 116^ Thus, the early inhibitory effect of BMS on PFKFB3 expression cannot be fully attributed to its inhibition of IKK signaling or p65 activation.^81, 82^ These previous studies demonstrating the relationship between IRF5 and p65 allow us to posit that there may be potential synergy or cooperation among these factors in regulating PFKFB3 expression. The importance of the finding that p65, STAT1 and IRF5 regulate PFKFB3 expression during immune response against Gram-negative bacteria/LPS may also extend to treatment of inflammatory diseases as these same transcription factors have been shown to drive polarization of macrophages into an M1 phenotype.^87, 88, 117, 118^ Collectively, our study uncovered that TBK1/IKKƐ/STAT1, IKK/p65, and IRF5 contribute to increased glycolysis through temporal upregulation of PFKFB3, which feeds back to positively regulate p65 expression, phosphorylation, as well as the activation of STAT1, thereby amplifying the inflammatory and ISG responses in macrophages responding to inflammatory stimuli.

### Limitations of the Study

Apart from p300, the role of other histone acetyl/lactyl transferases in p65 expression was not explored in this study. We observed a trend in which inhibiting p300 activity appeared to potently diminish p65 expression under normoxia more than under hypoxia, which may suggest that additional HATs regulate p65 expression, especially under hypoxia, and thus warrants further work. In addition, the interplay among the pathways we identified that regulate PFKFB3 expression requires further investigation to determine which regulator is most important and how that regulator affects p65 signaling and ISG responses. Lastly, we reported a global reduction in active histone PTMS upon glycolytic inhibition, which correlated with reduced expression of p65. However, we did not directly assess histone PTMs at the RELA loci, and whether this differs between normoxia and hypoxia during the course of LPS stimulation in human MDMs.

## MATERIALS AND METHODS

### Cell Culture

#### Generating Human Monocyte-Derived Macrophages (MDMs)

Peripheral blood mononuclear cells (PBMCs) were isolated using Ficoll-Paque Plus density centrifugation from buffy coat obtained from healthy donors from the New York Blood Center (NYBC). Briefly, monocytes were purified from isolated PBMCs via CD14-negative selection using a magnetic-based monocyte isolation kit. Cells were then resuspended with Roswell Park Memorial Institute (RPMI)-1640 supplemented with 10% Fetal Bovine Serum (FBS) and 1% penicillin/streptomycin and maintained at 37°C with 21% oxygen and 5% carbon dioxide. MCSF was added to RPMI at a final concentration of 25-50ng/ml. Cells were plated in 12- or 6-well plates at a given density. After 3-4 days of culture, cells were refreshed with RPMI supplemented with MCSF at the same concentration range and allowed to proceed until day 6-7, when they were fully differentiated into macrophages. 1×10^6^ cells were seeded for all assays except nuclear and cytoplasmic fractionation assays (2×10^6^ cells), and as otherwise stated in the assays. For hypoxic experimentation, cells were also first differentiated as described above. They were then maintained in a specialized hypoxic workstation (Whitley H35 HEPA hypoxy station) at 37°C with 1% oxygen, 5% carbon dioxide, and 94% nitrogen for the indicated time in each experiment. For the inhibitor-based experimental setup, cells were typically pretreated with compounds for 2h, then stimulated without LPS (mock) or with LPS (10 ng/ml) for 16h. After this, cells were harvested using cell scrapers and subjected to downstream assays.

#### Generating phorbol-12-myristate-13-acetate (PMA)-differentiated THP-1 macrophages from THP-1 monocytic cells

THP-1 monocytic cells were counted and resuspended with RPMI-1640 supplemented with 10% FBS and 1% penicillin/streptomycin and maintained at 37°C with 21% oxygen and 5% carbon dioxide, to which phorbol 12-myristate-13-acetate (PMA) was added at a final concentration of 16.2nM. Differentiation was allowed to proceed for 24 hours, after which the PMA-containing RPMI medium was removed, and cells were washed once with 1 mL PBS. After PBS removal, cells were replenished with fresh RPMI-1640 and allowed to rest for another 22-24 hours. The experimental setup with THP-1 macrophages follows the procedure described above for human MDMs.

### Western Blotting

Frozen cell pellets were lysed in 50 μL of Pierce^TM^ RIPA buffer supplemented with 10x phosphatase inhibitor, 100x protease inhibitor cocktail, and 2500x Universal nuclease. After at least 30 minutes of incubation on ice, the lysate was clarified by centrifugation at 16000 × g for 20 minutes at 4 °C. The supernatant was solubilized in 27 µL of 4× Laemmli and 10× DTT and denatured at 95 °C for 5 minutes. Lysates were loaded on 4 - 15% Mini-PROTEAN TGX gels (Bio-Rad) and transferred to a nitrocellulose membrane. Membranes were blocked with PBS Superblock for 1h and incubated with desired primary antibodies overnight on a rocker in a cold room maintained at 4 °C. This was followed by repeated washes of membranes with TBST (1X TBS + 0.1% Tween20), and their incubation for 1h at room temperature with HRP-linked secondary antibodies diluted in 2.5% blotto Milk in TBST. Membranes were repeatedly washed, and protein detection was achieved by Chemiluminescence using a ChemiDoc XRS+ Gel Imaging System (Bio-Rad). Image analysis was done using the Bio-Rad Image Lab software.

### Nuclear/Cytoplasmic Fractionation Assay

1-2 × 10^^6^ cells were plated in 6- or 12-well format. After successful differentiation, they were mock-treated or pretreated with inhibitors, then stimulated with LPS for 1h, 3h, or as indicated in the results section. The cells were scraped and harvested in cold PBS into 1.5 mL Eppendorf tubes, and pellets were obtained after centrifugation at 400 × g for 5 minutes. Cells were retained on ice. Cytoplasmic extraction was followed by using the Thermo Scientific NE-PER Nuclear and Cytoplasmic Extraction Reagents protocol. Briefly, cells were resuspended and vortexed in 100 or 200 μL of ice-cold CER I supplemented with 100× protease inhibitor, then incubated on ice for at least 10 minutes. After this, cold CER II was added to the tubes. Additional resuspension and vortexing were performed; the tubes were incubated on ice for at least 1 minute, and brief vortexing was repeated. The tubes were then centrifuged at 16000 × g for 5 minutes in a cold centrifuge (4 °C). The supernatant, representing the cytoplasmic fraction, was carefully transferred into newly labeled tubes, leaving the pellets behind. Both were stored in the - 20 °C. For nuclear extraction, the nuclear pellet was lysed in half the volume used for cytoplasmic extraction. Briefly, we employed Pierce^TM^ RIPA buffer supplemented with reagents described in the western blot section above, along with 100X DNase, to lyse pellets, which were then resuspended and vortexed vigorously. All other steps are as described for the Western blot above. Both cytoplasmic and nuclear lysates were solubilized in 4x Laemmli buffer and 10X DTT, then denatured at 95 °C for 5 minutes. Gel electrophoresis and protein detection were performed as previously described for Western Blotting.

### siRNA Transfection

Human MDMs were transfected with control or Silencer Select siRNA against IRF-1 and PFKFB3 for 16h using Lipofectamine RNAiMAX (Invitrogen) and OptiMEM. The media-containing transfection mix was replaced with fresh RPMI medium at 14h or 16h post-transfection. Cells were then treated with LPS for the indicated times. We confirmed siRNA knockdown of IRF-1 and PFKFB3 by western blot.

### RNA Extraction and quantitative RT-PCR

Total RNA was isolated from frozen cell pellets using the Zymo Quick RNA extraction kit, following the manufacturer’s instructions. cDNA was synthesized from 50 ng total extracted RNA using the reverse transcription system (Biorad) according to the manufacturer’s instructions. qRT-PCR was performed on the Biorad CFX96 Thermal cycler, using the SsoAdvanced Universal SYBR Green Supermix. GAPDH was used as an internal control. The following targets were amplified: REL A, PFKFB3, IRF-1, CCL5, CXCL10, TNF-α, and IL-6. Primer sequences were sourced from Integrated DNA Technologies (IDT), and information for each primer is listed in the key resource table below.

### Statistical Analysis

Statistical analyses and graphs were generated using GraphPad Prism. Statistical significance was determined using Student’s paired T-test or repeated measure One-Way ANOVA, and post-hoc was performed with Šídák’s multiple comparisons test; *p<0.05, **p<0.01, ***p<0.001, ****p<0.0001. Error bars indicate SDs.

## KEY RESOURCES TABLE

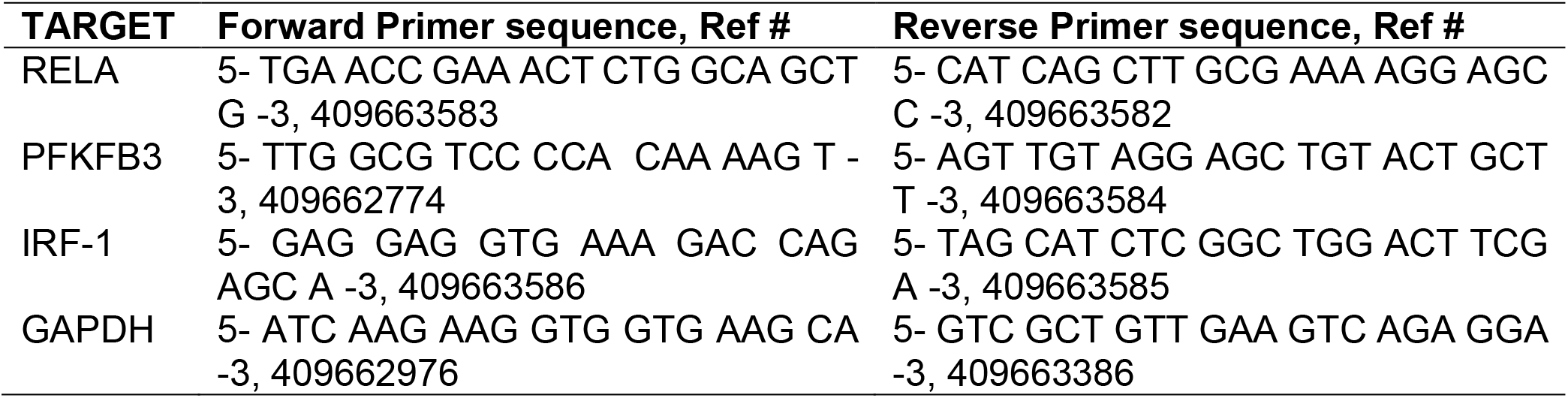
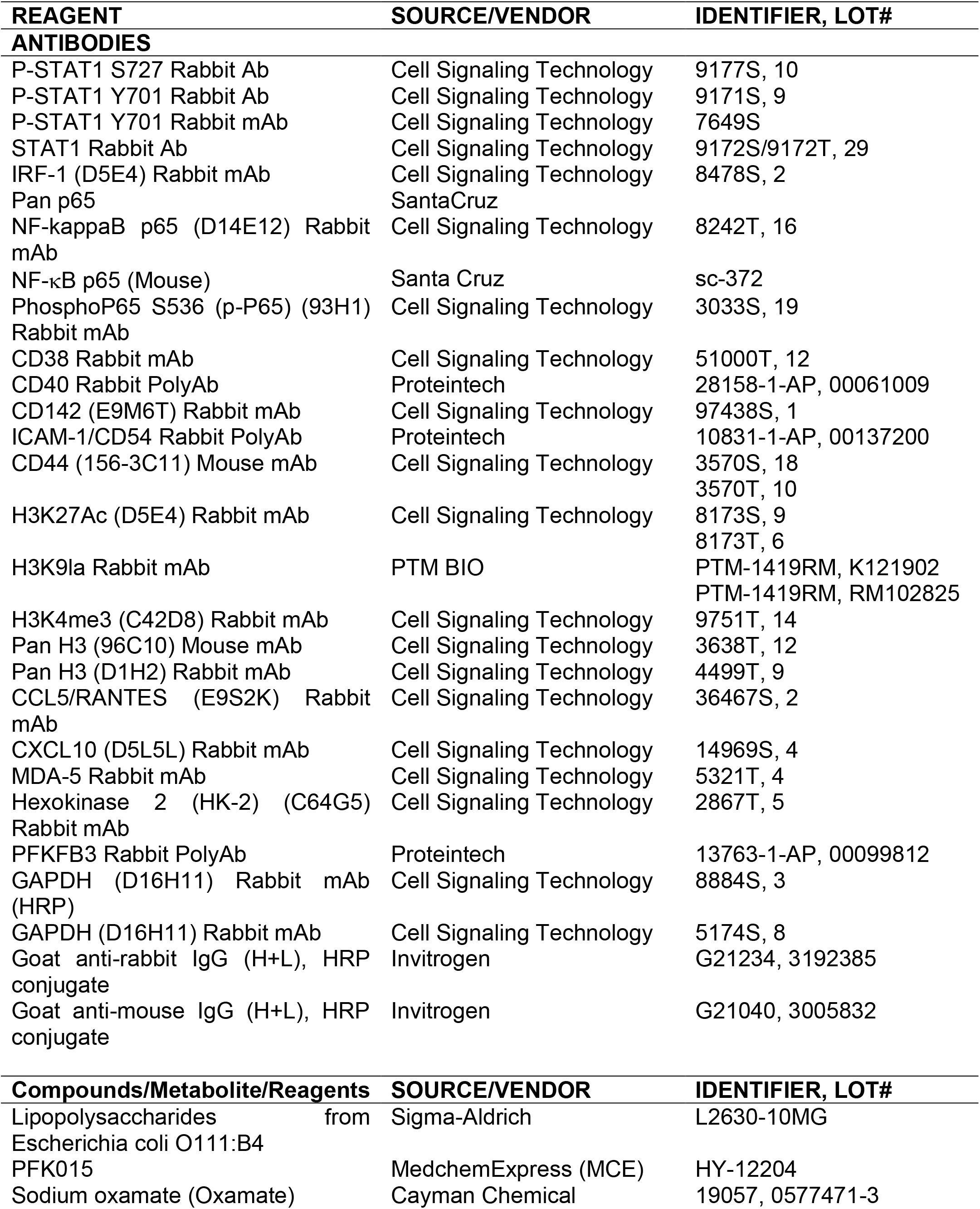
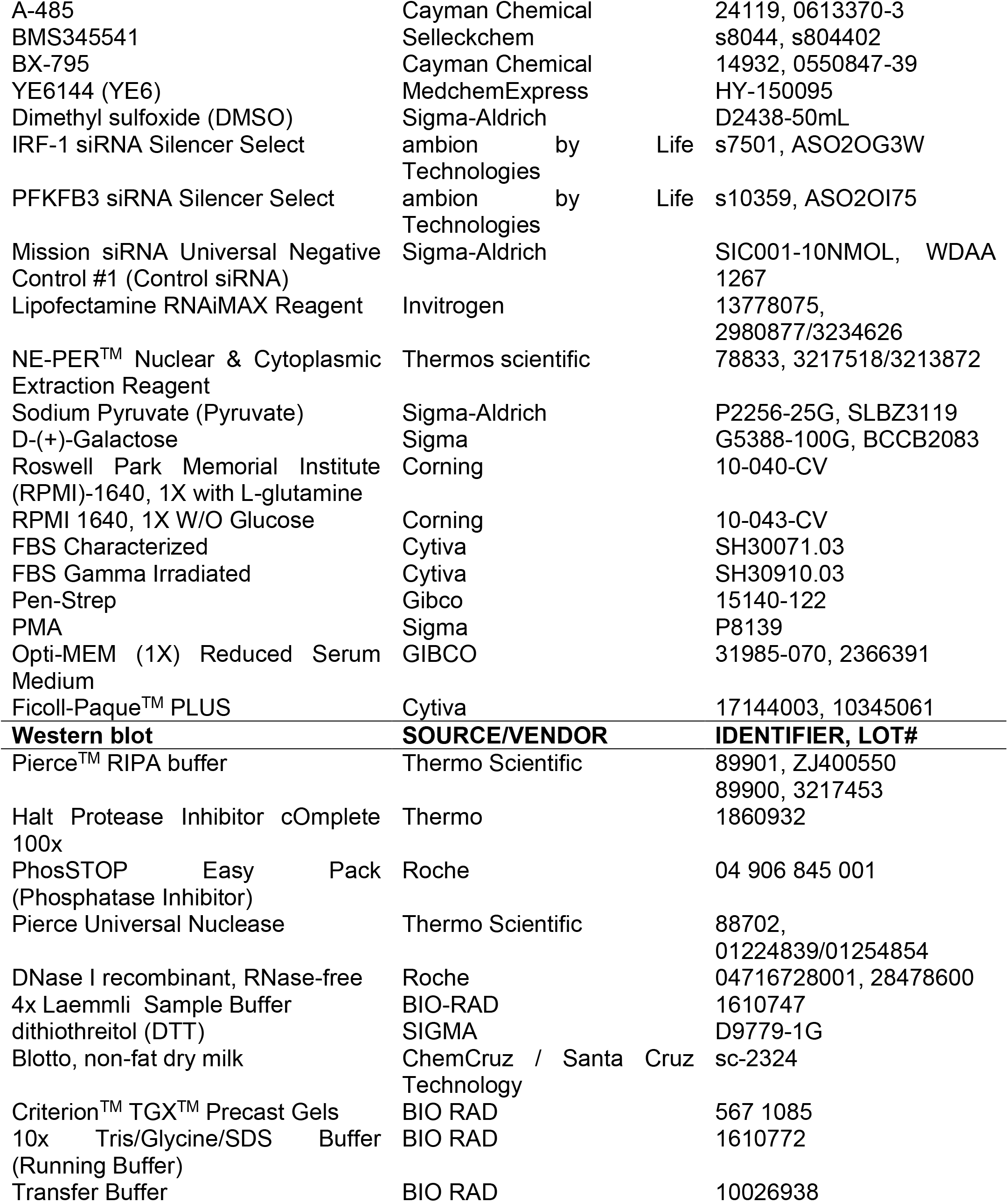
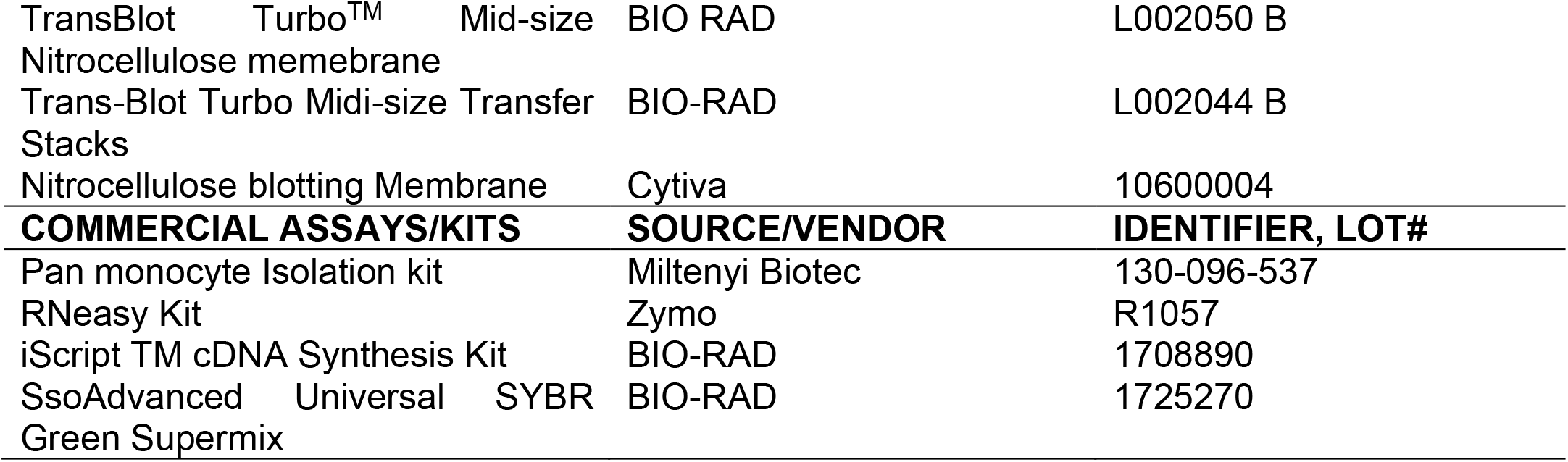

