## Supplementary Files for "Glucose Metabolism Mediates Feedback Control of Innate Immune Signaling in Human Macrophages"

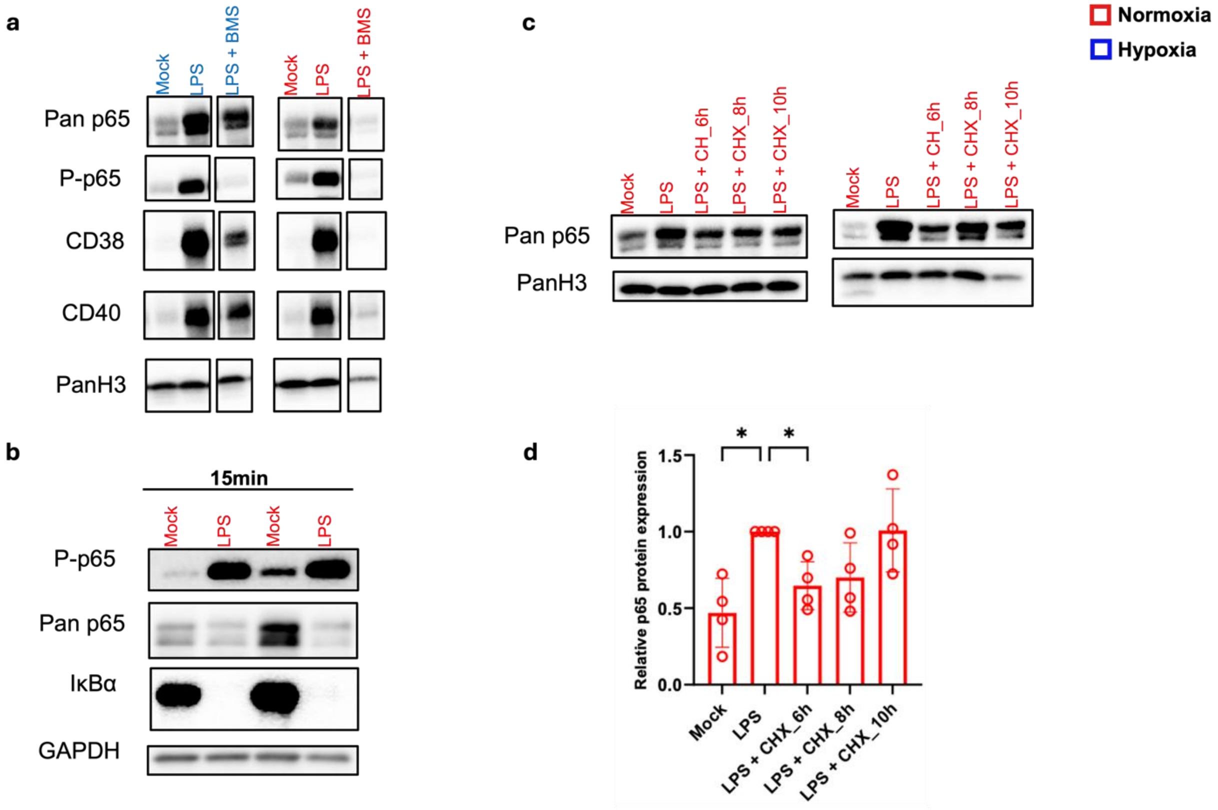


**Figure S1. Maximal p65 Protein Expression is due to Increased Protein Translation, and p65 persists in the nucleus at a Late Time Point**. (a) Primary human MDMs were pre-treated with the IKK complex inhibitor, BMS (2.5uM) for 2h, and then stimulated with LPS for 16h under normoxia (red) and hypoxia (blue). A representative blot of p65-target gene products, CD38 and CD40, as well as phospho and Pan p65 is shown. (b) Cells were untreated or stimulated with LPS for 15 min. Phospho p65 (P-p65), Ikappa B alpha (IκBα) and Pan p65 protein expression was detected by Immunoblot in 2 independent donors under normoxia. (c) Pan p65 expression was determined by immunoblot in cells stimulated with LPS for 16h, and to which cycloheximide (CHX, 25 μM) was added at 6h, 8h or 10h post-LPS stimulation. 2 independent donors are shown. (d) Densitometric analysis of Pan p65 in 4 independent donors is shown. Statistical significance was determined using repeated-measures one-way ANOVA and post hoc with Šídák's multiple comparisons test; *p<0.05, **p<0.01, ***p<0.001, ****p<0.0001. Error bars indicate SDs.

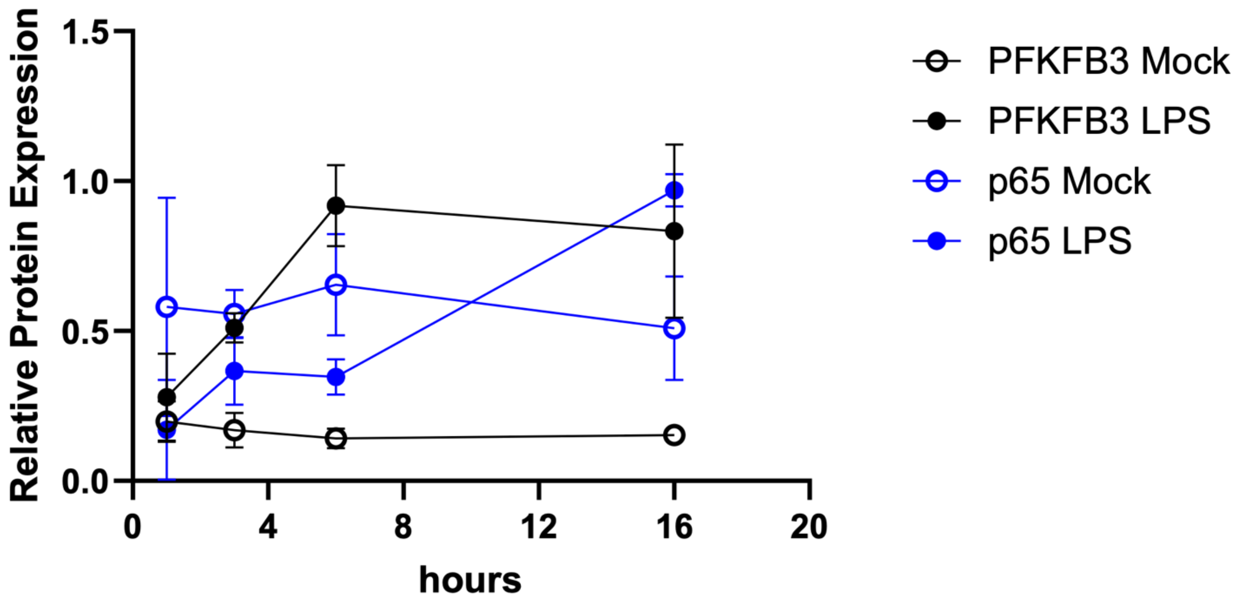


**Figure S2. PFKFB3 Expression is Increased Prior to Robust p65 Expression**. Primary human MDMs were treated with LPS for the 1,3,6 and 16h. Immunoblot and densitometric analysis were performed for p65 and PFKFB3. The graph represents a densitometric analysis of an overlay of PFKFB3 and p65 protein expression kinetics in stimulated cells under hypoxia.

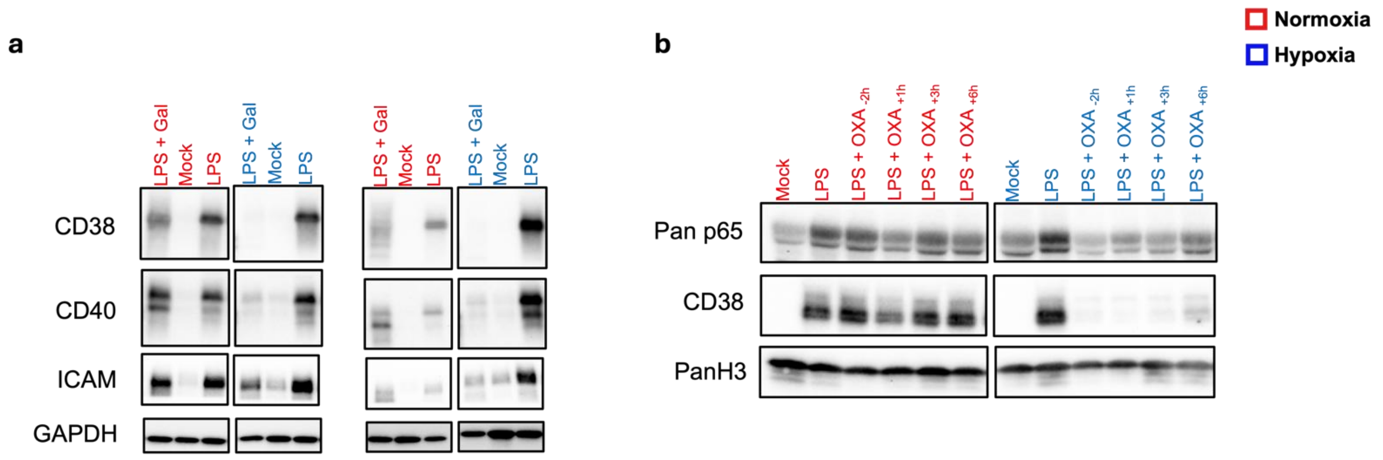


**Figure S3. Glucose Metabolism Regulates p65 and Late Protein Expression in LPS-Stimulated Human MDMs**. (a) Immunoblot of p65 target genes (CD38 and CD40) in 2 representative donors human MDMs cultured in galactose media prior to LPS stimulation for 16h. (b) Human MDMs were mock or pre-treated with OXA for 2h (-2h) or OXA was added either at 1h, 3h or 6h (represented as +) post LPS stimulation which lasted for 16h. Immunoblot result from a donor comparing p65 expression across different treatments.

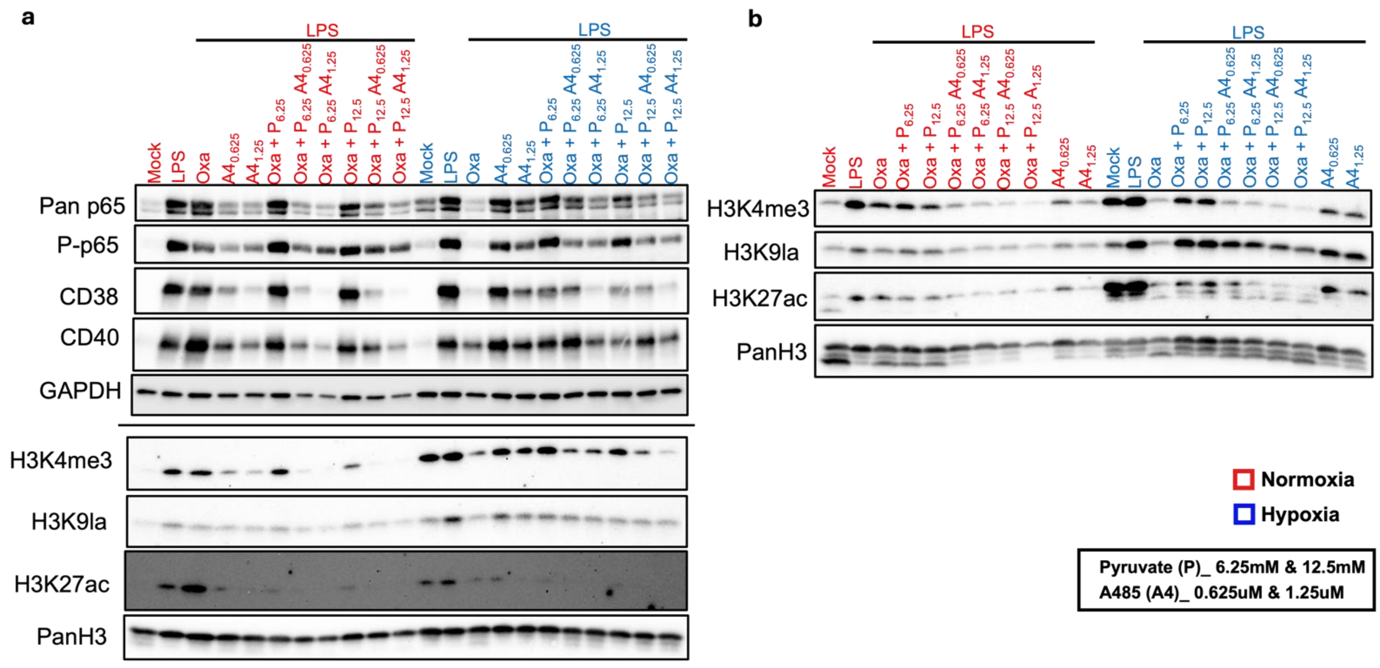


**Figure S4. p300 Activity Mediates p65 Expression and Rescue in Oxamate-treated Cells.** Human MDMs were mock- or pre-treated with 10 mM oxamate (OXA) or p300 inhibitor A485 (shortened as A4), respectively, at the indicated concentration for 2h. Afterwards, LPS and exogenous pyruvate (P) at the indicated concentrations were added to cells for 16h. (a) Immunoblotting was performed to detect Pan p65, P-p65, CD38, and CD40. GAPDH served as a loading control. (a, b) Immunoblot of histone PTMs H3K4me3, H3K27Ac, and H3K9La in 2 donors are shown. Pan H3 served as a loading control.

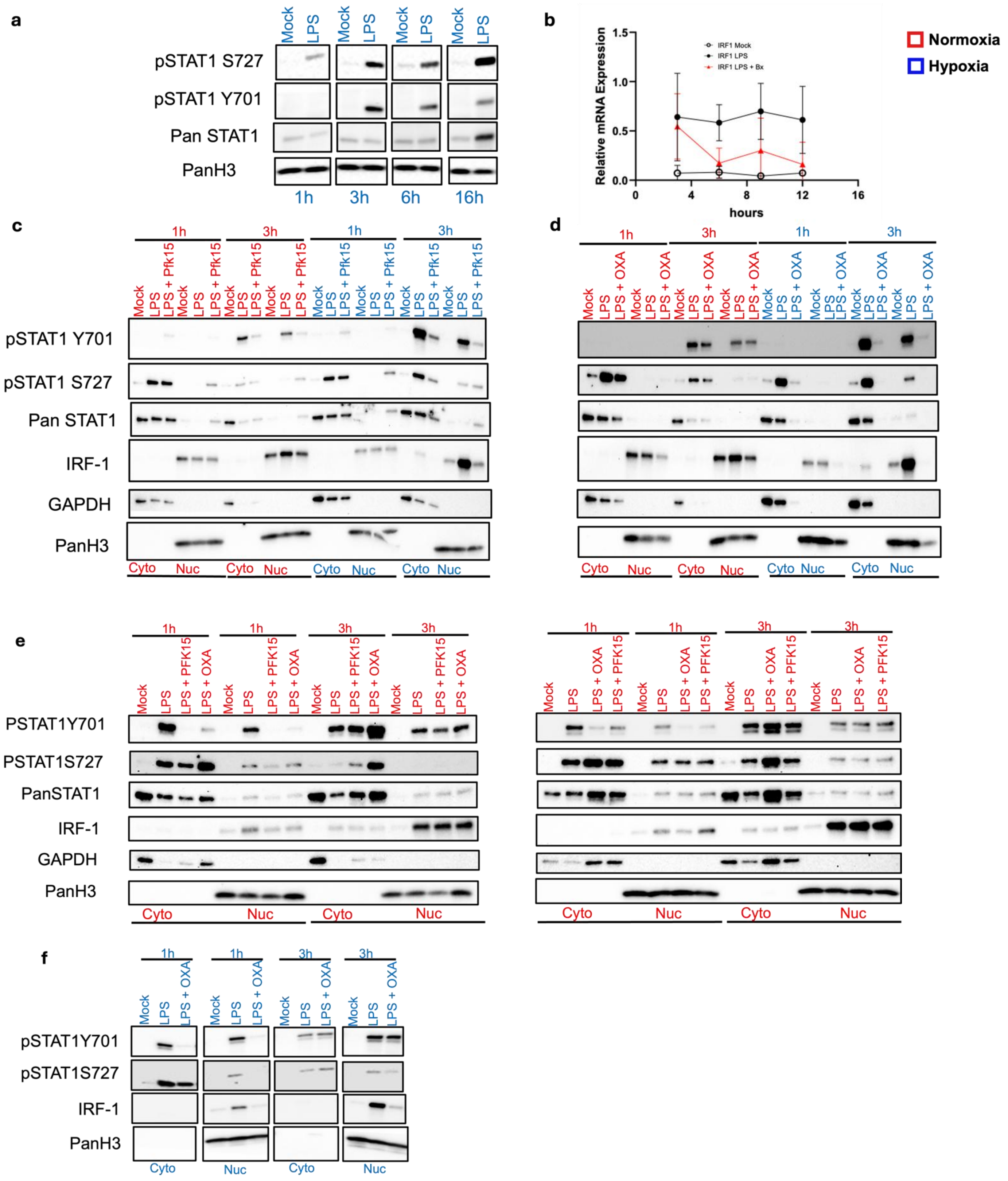
a
Mock
LPS
pSTAT1 S727
pSTAT1 Y701
Pan STAT1
PanH3
1h
3h
6h
16h
b
Relative mRNA Expression
hours
IRF1 Mock
IRF1 LPS
IRF1 LPS + Bx
Normoxia
Hypoxia
c
LPS + Pfk15
pSTAT1 Y701
pSTAT1 S727
Pan STAT1
IRF-1
GAPDH
PanH3
Cyto
Nuc
d
LPS + OXA
e
LPS + PFK15
PSTAT1Y701
PSTAT1S727
PanSTAT1
f
pSTAT1Y701
pSTAT1S727

**Figure S5. Glucose Metabolism is a Requirement for Early STAT-1 Y701 Phosphorylation and IRF-1 Expression**. (a) PMA-differentiated THP-1 macrophages were stimulated with LPS for the indicated time points under hypoxia, and an immunoblot for Pan STAT1, phosphorylated STAT1 at Y701 and S727 is shown. (b) qRT-PCR was performed on primary human MDMs pre-treated with BX795 and stimulated with LPS for the indicated times. The graph shows Relative IRF-1 mRNA expression, normalized to GAPDH. Statistical significance was determined using Student's paired T-test; $*p<0.05$, $**p<0.01$, $***p<0.005$. PMA-differentiated THP-1 macrophages were first pre-treated with (c) PFK15 or (d) Oxamate (OXA) and then stimulated with LPS for 1h and 3h respectively, under normoxia (red) and hypoxia (blue). Immunoblot for Pan STAT1, phosphorylated STAT1 at Y701 and S727, and IRF-1 protein was performed on total cell lysate. (e) Immunoblot analysis of nuclear fractionated primary human MDMs pretreated with OXA or PFK15 for 2h and stimulated with LPS for 1h and 3h respectively, under normoxia (red) and (f) hypoxia (blue). STAT-1 phosphorylation at Y701, S727, Pan STAT1, and IRF-1 is shown. GAPDH and PanH3 were used as controls for cytoplasmic and nuclear fractions, respectively. 2 independent donors are shown for normoxia, and 1 donor under hypoxia.

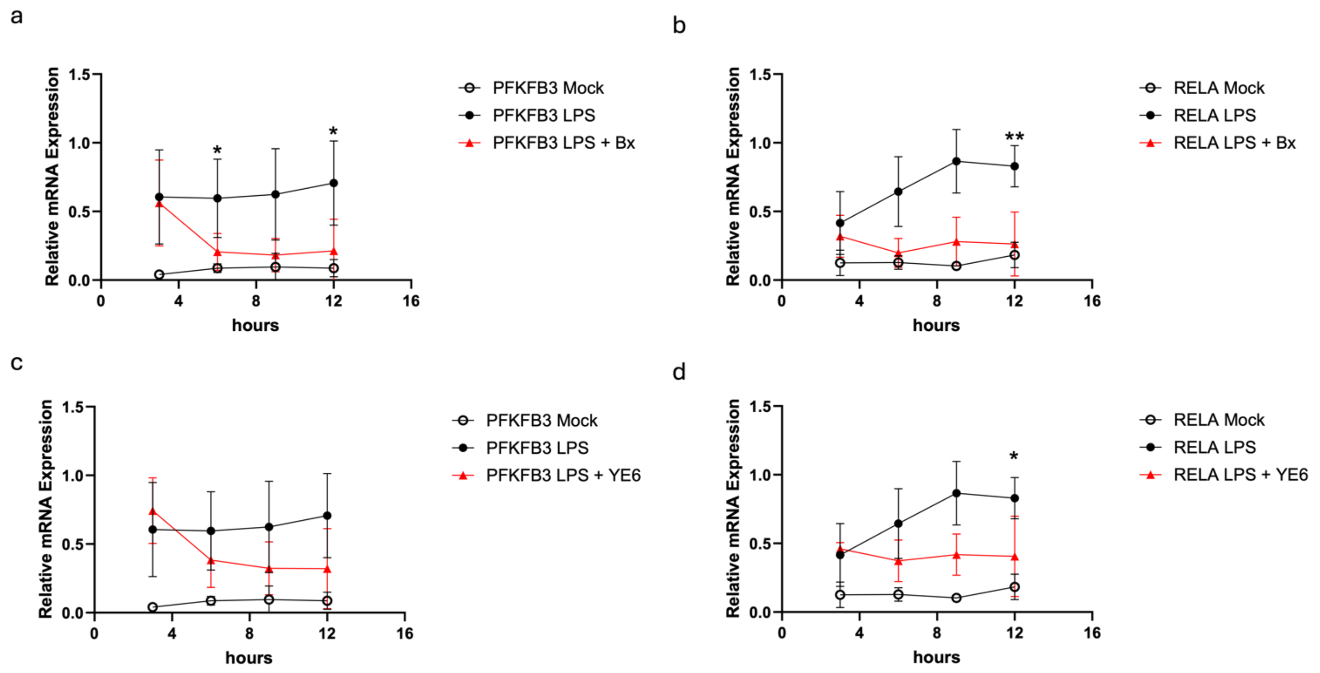


**Figure S6. Multiple TLR4-Driven Contribute to Temporal Induction of PFKFB3 in Primary Human MDMs**. Primary human MDMs were mock- or pre-treated with (a, b) BX795 (BX) and (c, d) IRF-5 inhibitor (YE6 - 1 μM) for 2h, and then stimulated with LPS for 3h, 6h, 9h, and 12h, respectively under normoxia. (a, b) Graphs depict relative mRNA expression of (a) PFKFB3 and (b) REL A/p65 in BX795-treated cells. (c, d) Graph depicts relative mRNA expression of (c) PFKFB3 and (d) Pan p65 in YE6-treated cells. Targets were normalized to GAPDH. Statistical significance was determined using Student's paired T-test; *$p<0.05$, **$p<0.01$, ***$p<0.001$, ****$p<0.0001$.
